# Human spinal cord organoids recapitulate developmental and disease-associated oligodendrocyte lineage signatures

**DOI:** 10.64898/2026.04.28.721458

**Authors:** Taylor Pio, Meghna Bettaiah, Ruizi Zhao, Supriya S. Wariyar, Sabra Mouhi, Emily J. Hill, Steven A. Sloan, Brain Organoid Hub, Jimena Andersen

## Abstract

Oligodendrocytes play essential roles in central nervous system development and homeostasis, and their dysfunction is a hallmark of numerous neurological disorders. However, human in vitro systems that support oligodendrocyte lineage progression while enabling the study of disease-relevant states remain limited. Here, we establish human spinal cord organoids (hSpO) and cortico-motor assembloids as platforms to model oligodendrocyte development, neuron-glia interactions, and cytokine-induced dysfunction. We show that hSpO generate oligodendrocyte lineage populations that transcriptionally resemble those found in the developing human spinal cord, and oligodendrocyte progenitor cells that exhibit physiologically-relevant functional properties, including migration and monosynaptic input from neurons. Exposure of assembloids to pro-inflammatory cytokines induces transcriptional changes across the oligodendrocyte lineage, characterized by altered lineage progression and acquisition of disease-associated gene expression programs that mirror signatures observed in multiple sclerosis patient tissue. Together, this work establishes hSpO and assembloids as in vitro systems for studying oligodendrocyte lineage development and disease-associated states in a human multi-cellular context.

## Introduction

Oligodendrocytes are essential regulators of central nervous system (CNS) function and homeostasis. They provide key contributions to neuronal metabolic support and ion regulation in addition to their well-established role in myelination and saltatory conduction. Oligodendrocytes arise from oligodendrocyte progenitor cells (OPCs), which exhibit diverse functional roles^1,2^, including synapse formation^3^, refinement^4^, and immune modulation^5,6^, and persist as the most abundant dividing cells in the adult CNS^7,8^. Both oligodendrocytes and OPCs have been shown to contribute to the onset and progression of neurological diseases like multiple sclerosis (MS)^9,10^, Alzheimer’s disease (AD)^11^, and amyotrophic lateral sclerosis (ALS)^12^. They can be among the earliest affected cells^13^, adopt inflammatory phenotypes^14^, and contribute to disease progression through both loss of supportive functions and acquisition of disease-like states. Despite these advances, the mechanisms underlying physiological and disease-associated states across the oligodendrocyte lineage remain poorly understood, reflecting limited access to experimentally tractable human systems capable of capturing early pathological changes beyond postmortem tissue.

We previously developed human spinal cord organoids (hSpO) from human induced pluripotent stem cells (hiPSCs) which recapitulate the neuronal diversity of the human spinal cord/hindbrain^15^. hSpO can be assembled with other region-specific organoids to create functionally integrated assembloids that model synaptic connections along the cortico-spinal-muscle axis^16^. These cortico-motor assembloids, which can be maintained in culture long-term, contain not only neurons, but also astrocytes and oligodendrocytes, and thus provide a platform to investigate cellular behaviors and interactions in complex neural environments. Detailed characterization of the oligodendrocyte lineage, their function, and their responses to insults within these assembloids, however, remain unexplored.

Here, we establish hSpO as a platform for studying human oligodendrocyte lineage biology. We show that hSpO and assembloid-derived oligodendrocyte lineage cells recapitulate human fetal spinal cord populations and exhibit key developmental behaviors, including migration into oligodendrocyte-naïve tissue and direct synaptic connections with spinal cord neurons. Furthermore, we establish an experimental paradigm to model oligodendrocyte disease-associated signatures. We show that a cytokine-induced pro-inflammatory environment drives a shift in oligodendrocyte state, characterized by altered lineage balance and acquisition of immune-associated gene expression that mirrors responses observed in primary human samples and resembles oligodendrocyte disease signatures in multiple sclerosis tissue.

## Results

### Oligodendrocyte lineage in hSpO

To study the oligodendrocyte lineage in hSpO, we first performed bulk RNA sequencing on organoids derived from three hiPSC lines from day (d) 75 to d200 in culture (**Figures 1A** and **S1A**; **Table S1A** lists hiPSC lines used in various experiments). Analysis indicated that *SOX10, PDGFRA*, and *MAG*, markers specific to the oligodendrocyte lineage, OPCs, and myelin-forming/mature oligodendrocytes (MFOL/MOL) respectively, increased over time and plateaued by d120 (**Figure 1B**). Immunohistochemistry in whole hSpO revealed abundant SOX10+ cells, as well as OPCs (PDGFRA+) and more mature oligodendrocytes (MBP+) at these timepoints (**Figures 1C** and **1D**; antibodies listed in **Table S1C**). Neurons (NFH+) and astrocytes (GFAP+) were also distributed throughout the organoids and in close proximity to cells of the oligodendrocyte lineage (GALC+) at d120 (**Figures S1B** and **S1C**).

**Figure 1.**
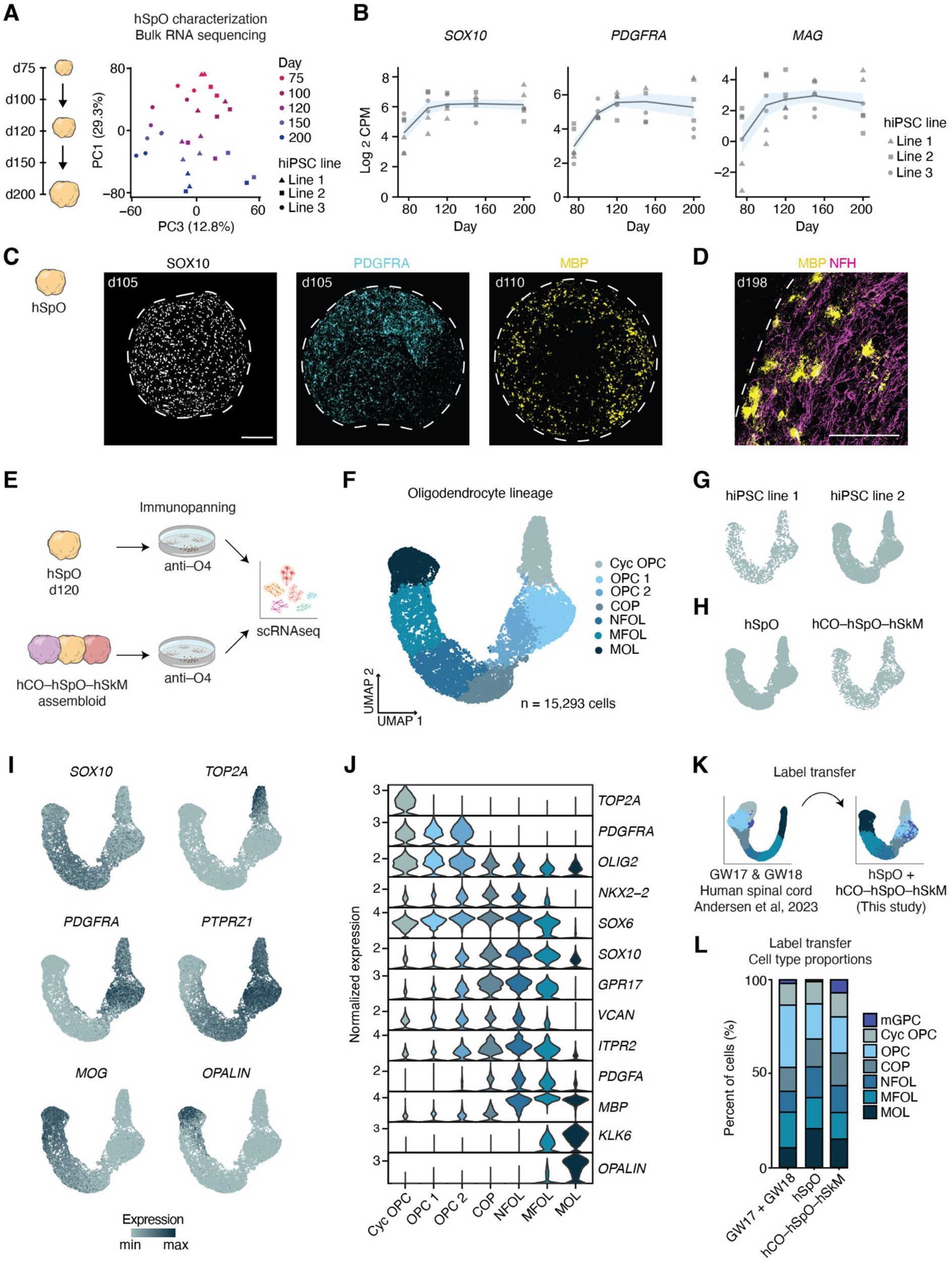
Oligodendrocyte lineage in hiPSC-derived hSpO and assembloids. **(A)** Principal Component Analysis (PCA) plot with hSpO samples over time showing PC1 and PC3 and colored by age. n = 2 replicates shown per hiPSC line and timepoint, with 3 hiPSC lines (represented with shapes; triangle, square, and circle) and 4 timepoints. Each replicate consists of 2-3 pooled organoids. See **Figure S1A** for plot showing PC1 and PC2. **(B)** Log_2_ Counts Per Million (CPM) expression of canonical oligodendrocyte lineage markers from day 75 to day 200. n = 2 replicates per hiPSC line and timepoint, with 3 hiPSC lines and 4 timepoints. Each replicate consists of 2-3 pooled organoids. See **Figure S1B** for expression of neuron and astrocyte markers. **(C)** Representative images of oligodendrocyte lineage markers SOX10, PDGFRA, and MBP in hSpO whole organoid immunohistochemistry at days 105–110. **(D)** Representative image of oligodendrocyte (MBP) and neuron (NFH) immunohistochemistry in hSpO cryosection at day 198. **(E)** Schematic detailing hSpO and hCO–hSpO–hSkM culture. hSpO was used at day 120 in both conditions (hCO–hSpO–hSkM 10 weeks of assembly). Oligodendrocyte lineage cells were enriched through immunopanning and captured for single cell RNA sequencing (scRNA seq, 10x Chromium). **(F)** UMAP visualization of single cell gene expression of organoid-derived oligodendrocyte lineage cells colored by cell type (n = 15,293 cells from 2 hiPSC lines). See **Figure S1D** for quality control metrics. **(G)** UMAP plot separated by the hiPSC line they were derived from. **(H)** UMAP plot separated by the culture condition they were derived from. **(I)** Feature plots showing gene expression of selected oligodendrocyte lineage marker genes. Color scale represents log-transformed normalized gene expression (counts per 10,000), scaled to the minimum and maximum values across all cells shown per gene. **(J)** Violin plots showing gene expression of selected oligodendrocyte lineage marker genes in each cell cluster. Y-axis represents normalized gene expression, log transformed (counts per 10,000). **(K)** Schematic showing cell type label transfer from primary human spinal cord oligodendrocyte lineage dataset at Gestation Weeks (GW) 17 and 18 onto organoid-derived oligodendrocyte lineage dataset (Andersen, Thom et al, 2023 and this study, respectively). See **Figures S1E** and **S1F** for primary dataset. **(L)** Stacked bar plot showing the proportion of cell types along the oligodendrocyte lineage based on the label transfer annotations from the primary dataset (conditions are hSpO, hCO–hSpO–hSkM, or primary dataset). See **Figure S1G** for a comparison of the label transfer annotation with our manual Seurat cluster annotation. Scale bar: 200 μm (**C**) and 100 μm (**D**).

To more comprehensively resolve specific stages along the oligodendrocyte lineage, we performed single cell RNA sequencing (10x Chromium). Our previous work described the presence of MBP+ oligodendrocytes in 10-week cortico-motor assembloids^15^. To establish whether assembly is required for oligodendrocyte lineage progression, we included both hSpO alone and cortical-spinal-muscle assembloids (hCO-hSpO-hSkM, 10 weeks after assembly) in our analysis, with hSpO at d120 in both conditions. We used immunopanning against the OPC/oligodendrocyte-specific marker O4 to enrich for cells in the lineage prior to capture and sequencing^17^ (**Figure 1E**). After quality control and filtering, we obtained transcriptomes for 15,293 cells across the two conditions and in two hiPSC lines. We identified seven high-quality clusters and annotated them using canonical markers previously reported across the oligodendrocyte lineage, including *PDGFRA, NKX2-2, ITPR2, MOG*, and *KLK6*^18,19^ (**Figures 1I, 1J** and **S1D**). These clusters consisted of cycling OPCs (Cyc OPC), OPC1, OPC2, committed oligodendrocyte precursors (COP), newly formed oligodendrocytes oligodendrocytes (NFOL), (MFOL), myelin-forming and mature oligodendrocytes (MOL; **Figure 1F**; highly expressed markers in each cluster listed in **Table S1G**). Oligodendrocyte populations were similar across two hiPSC lines and between hSpO and assembloid conditions (**Figures 1G** and **1H**; differentially expressed markers between conditions listed in **Table S1H**), suggesting that assembly does not influence overall lineage progression. To validate the identity of these populations, we performed label transfer using our previously generated human fetal spinal cord dataset from gestational weeks (GW) 17 and 18 (**Figures 1K** and **S1E–G**) as a reference^19^. This transferred reference-based annotation aligned with our marker-based identification, corroborating the cell type identity of these populations.

Prior studies, including our own, have reported the presence of multipotent glial progenitors (mGPCs^19,20^, also described as giPCs^21^, pre-OPCs^22^, pri-OPCs^23^, and bMIPCs^24^) within the developing human nervous system. Though mGPCs did not initially form their own cluster in our hSpO/assembloid dataset, a group of cells characterized by the markers *EGFR, PDGFRA*, and *AC007402*.*1* was labeled as mGPC after label transfer (**Figures S1G** and **S1H**). Analysis of the cell type composition between organoids, assembloids, and primary tissue indicated overall comparable proportions, with hCO-hSpO-hSkM exhibiting an increase in mGPCs compared to hSpO alone (**Figure 1L**). While standard dorsal forebrain (cortical) organoid protocols typically do not generate oligodendrocyte lineage cells without additional patterning molecules^25–27^, the presence of mGPC signatures has been observed^28^, pointing at hCO as a possible source for these cells in our dataset.

### Functional characterization of oligodendrocyte lineage cells in hSpO

We next investigated whether hSpO-derived oligodendrocyte lineage cells displayed functional features associated with their roles in the CNS. During nervous system development, OPCs are generated in discrete zones and subsequently migrate to populate their target regions. To test migratory capacity, we infected hSpO with a lentivirus expressing GFP under the MCS5 enhancer element of the SOX10 gene^29^ (LV-SOX10-GFP), directing GFP expression to oligodendrocyte lineage cells and enabling their long-term tracking. We then assembled hSpO with oligodendrocyte-naive hCO at d95 and tracked migration of GFP cells (**Figure 2A**; all viruses listed in **Table S1D**). Quantification of area of GFP+ cells in hCO normalized to the initial area of GFP in hSpO indicated that over the course of two weeks, SOX10-GFP cells populated the hCO, demonstrating robust migratory capacity (**Figures 2B** and **2C**). A unique feature of OPCs is their ability to form bona fide synapses with neurons^30^.

**Figure 2.**
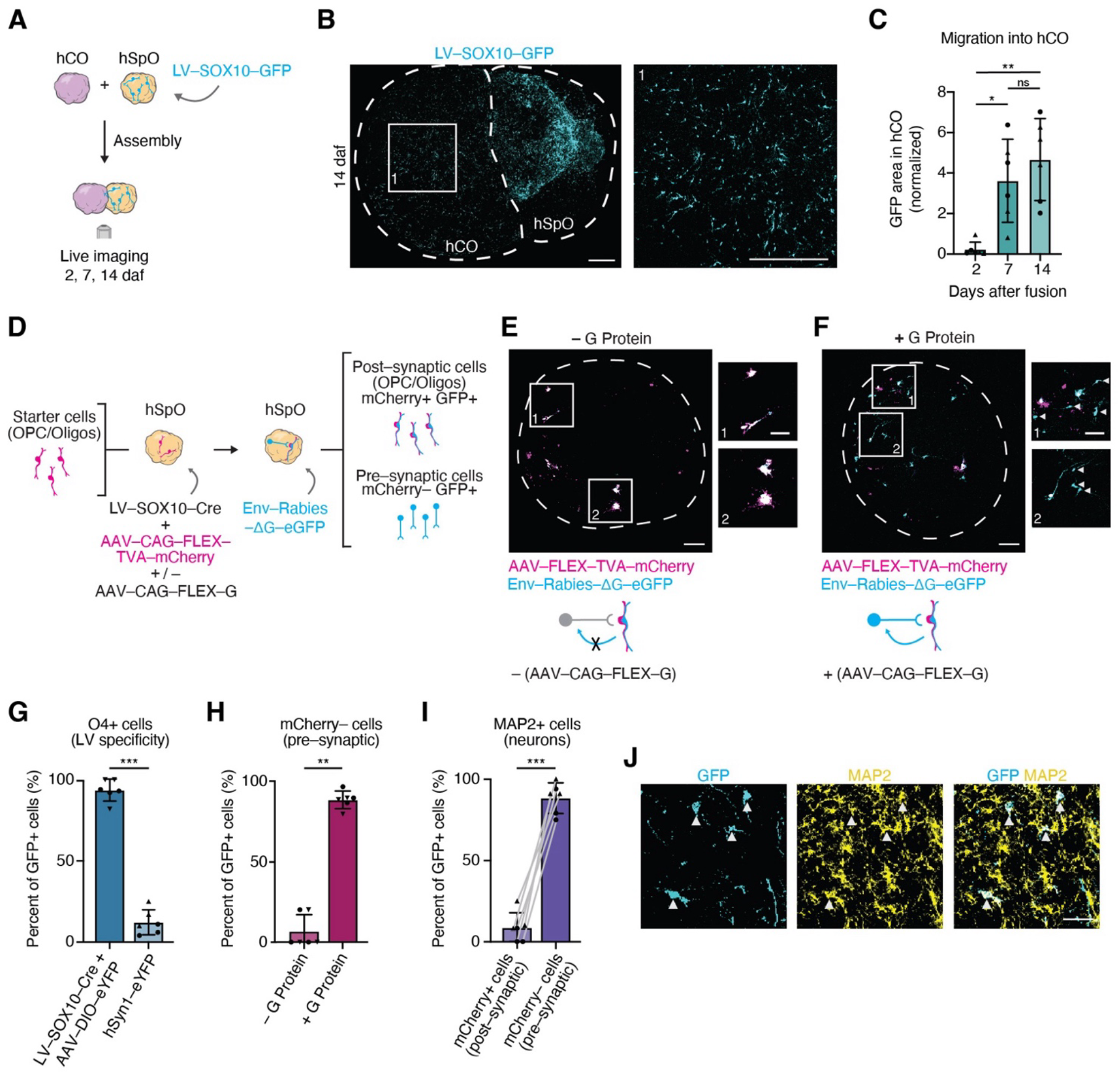
Functional integration of oligodendrocyte lineage cells in hiPSC-derived organoids. **(A)** Schematic detailing hSpO infection with LV-SOX10-GFP, assembly with hCO, and imaging at 2, 7, and 14 days after fusion (daf). **(B)** Representative image of live hCO–hSpO assembloid at 14 daf. Inset showing GFP+ cells migrated into hCO. **(C)** Quantification of GFP+ area in hCO normalized to starting GFP+ area in hSpO at 2 daf. n = 3 assembloids per line from 2 hiPSC lines. Mean ± SEM: d2 = 0.23 ± 0.15, d7 = 3.62 ± 0.84, d14 = 4.67 ± 0.83; Kruskal-Wallis test with Dunn’s multiple comparisons: d2 vs d7 p = 0.028, d2 vs d14 p = 0.006, d7 vs d14 p > 0.999. Statistical significance is denoted as p < 0.05 (*) and p < 0.01 (**). **(D)** Schematic detailing monosynaptic retrograde viral tracing experiment specific to oligodendrocyte lineage starter cells. hSpO are infected with LV-SOX10-Cre for oligodendrocyte lineage-specific Cre-recombinase expression, AAVDJ-FLEX-TVA-mCherry for Cre-dependent rabies entry, and AAVDJ-FLEX-G for Cre-dependent monosynaptic spread. After two weeks, hSpO are infected with enveloped G-deleted rabies-eGFP (EnV-Rabies-ΔG-eGFP). Post-synaptic (starter cells, SOX10+ oligodendrocyte lineage cells) are identified as mCherry+GFP+ cells. Pre-synaptic cells are identified as mCherry–GFP+. (**E, F**) Live imaging of hSpO infected with AAV-FLEX-TVA-mCherry and enveloped G-deleted rabies-eGFP in the absence (**E**) or presence (**F**) of G protein. Arrows in insets (**F**) show pre-synaptic rabies-eGFP+ cells that are mCherry–. **(G)** Quantification of O4+ cells colocalizing with eYFP in hSpO infected with either LV-SOX10-Cre and AAV-DIO-eYFP or only AAV-hSyn1-eYFP. Mean ± SEM: SOX10-Cre = 93.5% ± 2.7%, hSyn1-eYFP = 12.2% ± 3.2%; n = 6 organoids from 2 hiPSC lines, with 2 sections quantified and averaged per organoid; one-way ANOVA with Tukey’s multiple comparisons: SOX10-Cre vs hSyn1-eYFP p<0.0001. LV-SOX10-Cre was used at a concentration of 1:5,000, see **Figures S2A** and **S2B** for comparison of virus specificity at differing concentrations. Statistical significance is denoted as p < 0.001 (***). **(H)** Quantification of the percentage of GFP+ cells that are pre-synaptic cells (rabies-eGFP+ cells that are mCherry–) in organoids with or without G protein. Mean ± SEM: –G protein = 6.7% ± 4.2%, +G protein = 87.8% ± 2.2%; n = 6 organoids per condition from 2 hiPSC lines, with 1–3 sections quantified and averaged per organoid; Mann-Whitney test, p = 0.002. Statistical significance is denoted as p < 0.01 (**). **(I)** Quantification of the percentage of rabies-eGFP+ cells that co-express the neuronal marker MAP2 in either post-synaptic cells (mCherry+) or pre-synaptic cells (mCherry–). Mean ± SEM: GFP+ mCherry+ = 8.6% ± 3.7%%, GFP+ mCherry– = 88.3% ± 3.9%; n = 6 organoids per condition from 2 hiPSC lines, with 1–3 sections quantified and averaged per organoid; lines connecting data points from the same organoid; paired t test, p < 0.0001. Statistical significance is denoted as p < 0.001 (***). **(J)** Representative immunohistochemistry image showing expression of rabies-derived GFP (pre-synaptic cells) colocalized with MAP2 (neurons). Arrows show pre-synaptic rabies-eGFP+ cells that are MAP2+. Scale bar: 500 μm (**B**), 200 μm (**E, F**), 100 μm (insets in **E, F**), 50 μm (**J**).

To test whether hSpO-derived OPCs receive direct synaptic input, we used a modified rabies virus tracing approach that specifically labels cells that are monosynaptically connected to oligodendrocyte lineage cells. This system ensures specificity in two steps: first, a modified rabies virus pseudotyped with EnvA and expressing a GFP reporter can only infect cells that express the avian TVA receptor; second, the virus can only spread retrogradely across synapses to connected neurons if the starter cells also express rabies glycoprotein (G protein), enabling visualization of the presynaptic partners.

To restrict starter cells to the oligodendrocyte lineage, we expressed Cre recombinase in oligodendrocyte lineage cells using a SOX10-specific lentiviral vector (LV-SOX10-Cre). Cre-dependent AAVs encoding TVA with mCherry (AAV-FLEX-TVA-mCherry) and G protein (AAV-FLEX-G) were co-delivered, such that only Cre-expressing oligodendrocyte-lineage cells expressed TVA-mCherry and G protein and were competent for rabies infection and monosynaptic spread (**Figure 2D**). Infection of hSpO with an EnvA-pseudotyped, G-deleted rabies virus expressing GFP (EnvA-ΔG-rabies-eGFP) two weeks later resulted in GFP expression in SOX10+ starter cells (mCherry+GFP+; post-synaptic cells) and retrograde labeling of directly connected pre-synaptic cells (mCherry–GFP+; **Figures 2F**).

We first validated the lineage specificity of LV-SOX10-Cre by co-infecting hSpO with a Cre-dependent eYFP reporter under a universal mammalian promoter (AAV-CAG-DIO-eYFP; **Figure S2A**). Immunohistochemistry revealed that approximately 95% of eYFP+ cells co-expressed the oligodendrocyte marker O4, confirming selective targeting of the oligodendrocyte lineage (**Figures 2G, S2B** and **S2C**). We next confirmed that EnvA-ΔG -rabies-eGFP virus entry was strictly dependent on TVA expression, as no GFP signal was detected in the absence of TVA (**Figure S2D** and **S2E**).

To verify that labeling of presynaptic cells required G protein-mediated trans-synaptic spread, we compared conditions with and without G protein delivery. In the absence of G protein, GFP labeling was largely restricted to starter cells (mCherry+GFP+), with minimal presence of connected cells expressing only GFP (mCherry– GFP+). Delivery of G protein resulted in a 13-fold increase in mCherry–GFP+ cells, indicating robust retrograde labeling of presynaptic cells (6.7% −G versus 87.8% +G; **Figures 2E, 2F** and **2H**). Immunohistochemistry of hSpO in cryosections revealed that approximately 90% of these mCherry–GFP+ presynaptic cells co-expressed the neuronal marker MAP2 (**Figure 2I** and **2J**), indicating that neurons form monosynaptic connections onto oligodendrocyte lineage cells in hSpO.

### Modeling neuroinflammation-induced lineage disruption

OPC and oligodendrocyte dysfunction is observed across neurological diseases including MS^31^, AD^32^, ALS^33^, and schizophrenia^34^, characterized by loss of supportive functions, disrupted lineage, and acquisition of disease signatures. A common feature in these conditions is neuroinflammation, often accompanied by signaling from activated microglia. Organoids patterned towards neuroectoderm lack microglia and immune-like cells, however previous work has demonstrated that treatment with microglial-derived cytokines TNF-α, IL-1α, and C1q (TIC) can induce astrocyte reactivity in mouse models^35^, human cortical tissue, and human organoid models^36,37^. Given increasing evidence for cytokine-responsive, immune-competent oligodendrocyte lineage cells^38^, and the extensive cross-talk between astrocytes and oligodendrocytes^39^, we investigated if TIC treatment could induce a reactive, disease-like state in the hSpO oligodendrocyte lineage. We treated hCO-hSpO-hSkM assembloids with TIC for eight weeks, then immunopanned for O4+ cells and performed single-cell RNA sequencing (**Figure 3A**). As reactive astrocytes are a hallmark of neuroinflammation, we also immunopanned astrocytes from 8-week TIC-treated assembloids for single-cell RNA sequencing separately. We found that TIC-treated astrocytes indeed displayed a reactive signature^35^ and upregulation of known reactive genes including *C3, LCN2*, and *GFAP*, along with the chitinase-like inflammatory gene *CHI3L1* and the Alzheimer’s disease risk factor *APOE* (**Figures S3A, S3B, S3D and S3H-S3K**; differential expression data for astrocyte treatment listed in **Table S1K**).

**Figure 3.**
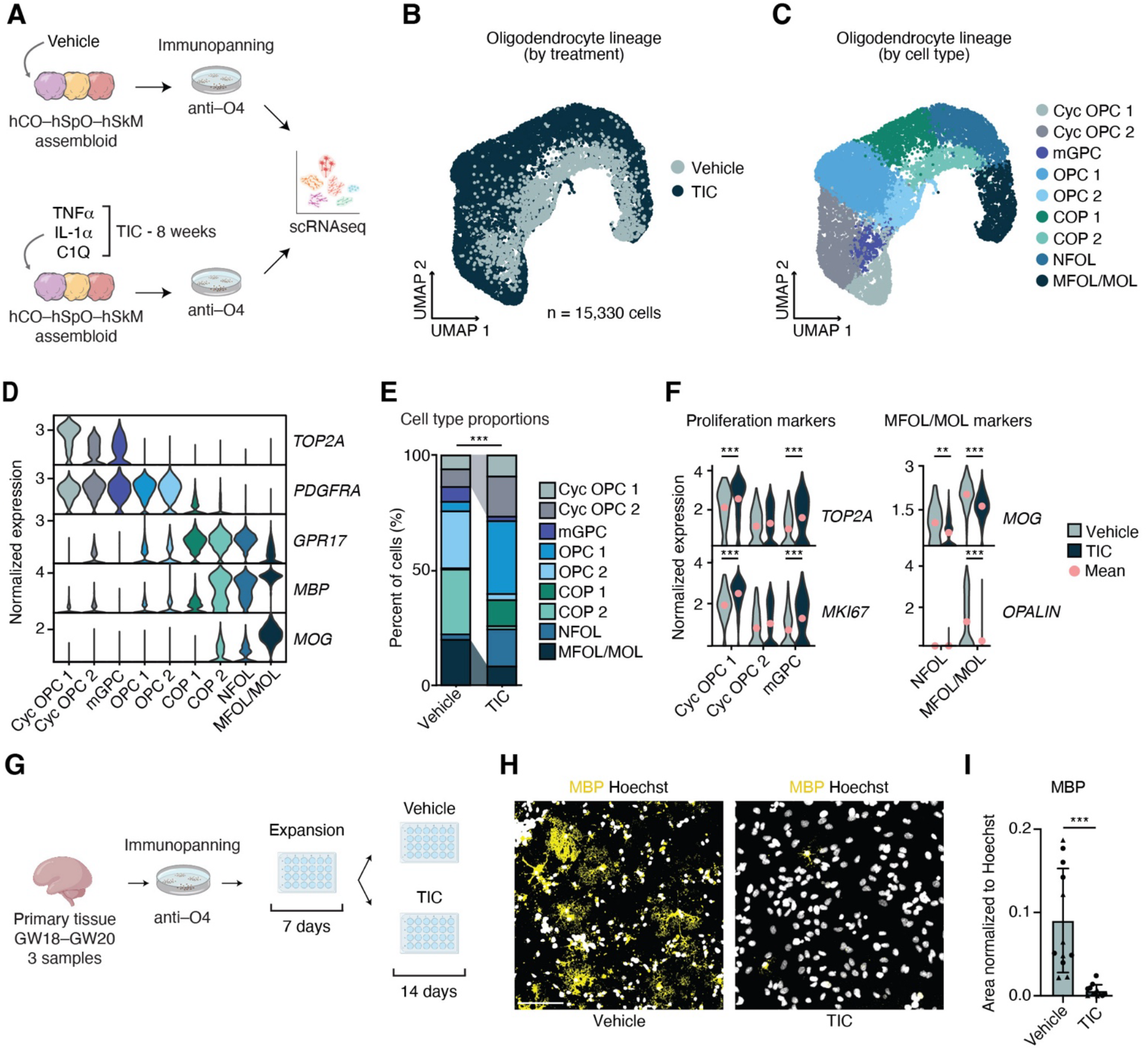
Cytokine-induced oligodendrocyte lineage disruption. **(A)** Schematic detailing assembloid cultures treated with vehicle or TNF–α, IL–1α, and C1q (TIC) for 8 weeks (hSpO day 120, hCO–hSpO–hSkM 10 weeks of assembly). Oligodendrocyte lineage cells were enriched through O4 immunopanning and captured for single cell RNA sequencing (scRNAseq, 10x Chromium). Astrocytes were also immunopanned using HEPACAM and captured separately. **Figure S3A** shows panning scheme, and **Figure S3B** shows UMAP with both populations. **(B)** UMAP plot showing a subcluster of the oligodendrocyte lineage with cells grouped by treatment (vehicle or TIC). See **Figure S3F** for cells grouped by hiPSC line. **(C)** UMAP plot showing the organoid-derived oligodendrocyte lineage, with cells grouped by cell type (annotated Seurat clusters; n = 15,330 cells from 2 hiPSC lines). **(D)** Violin plots showing gene expression of selected oligodendrocyte lineage marker genes in each annotated cluster (clusters not split by treatment). Y-axis represents normalized gene expression, log transformed (counts per 10,000). **(E)** Stacked bar plot showing the percentage of cells in each of the clusters separated by treatment (vehicle or TIC). See **Figure S3G** for the percentage of cells in each treatment group separated by cluster. Chi-squared analysis of cell type proportion changes, p <0.0001. Statistical significance is denoted as p < 0.001 (***). **(F)** Violin plots showing proliferation (*TOP2A, MKI67*) and more mature (MFOL/MOL) oligodendrocyte (*MOG, OPALIN*) gene expression in vehicle or TIC treatment groups. Y-axis represents normalized gene expression, log transformed (counts per 10,000). Wilcoxon rank sum test; Cyc OPC 1: *TOP2A* p < 0.0001, *MKI67* p < 0.0001; Cyc OPC 2: *TOP2A* p = 1, *MKI67* p = 1; mGPC: *TOP2A* p = 0.0001, *MKI67* p-value = 0.0001; NFOL: *MOG* p = 0.007, *OPALIN* p N/A; MFOL/MOL: *MOG* p <0.0001, *OPALIN* p <0.0001. Statistical significance is denoted as p < 0.05 (*), p < 0.01 (**), and p < 0.001 (***). **(G)** Schematic detailing enrichment of oligodendrocyte lineage cells from fetal cortical tissue via immunopanning. Oligodendrocyte-enriched cultures underwent 7 days of expansion, then were treated with either vehicle or TIC for 14 days. See **Figure S4A–C** for validation of oligodendrocyte enrichment. **(H)** Representative images showing immunohistochemistry of oligodendrocytes (MBP) and nuclei (Hoechst) in vehicle and TIC-treated oligodendrocyte-enriched cultures. **(I)** Quantification of MBP area normalized to Hoechst area in vehicle or TIC-treated oligodendrocyte-enriched cultures. Mean ± SEM: Vehicle = 0.090 ± 0.018, TIC = 0.0061 ± 0.0021; n = 3 samples GW18– GW20, 2 wells per sample and 2 images per well; Wilcoxon test, p = 0.0005. Statistical significance is denoted as p < 0.001 (***). Scale bar: 100 μm (**H**).

We then analyzed 15,330 oligodendrocyte lineage cells from two hiPSC lines and annotated clusters using oligodendrocyte lineage markers as before (**Figures 3B–D** and **S3C–F;** highly expressed markers in each cluster listed in **Table S1I**). Several clusters, including cycling (Cyc) OPC1, Cyc OPC2, OPC1, COP1, and NFOL, were predominantly composed of TIC-treated cells, with minimal representation from vehicle-treated cells (**Figures 3B, 3C** and **S3G**), demonstrating a transcriptional shift across the lineage with cytokine exposure. Cell type proportion analysis revealed that TIC treatment increased representation of proliferative clusters and decreased the proportion of the most mature MFOL/MOL cluster (**Figure 3E**). At the gene expression level, TIC-treated samples also showed elevated expression of proliferation markers *TOP2A* and *MKI67* and reduced expression of MFOL/MOL markers *MOG* and *OPALIN* (**Figure 3F**).

To determine whether these phenotypes reflect biologically relevant responses to TIC treatment, we obtained human fetal cortical tissue spanning GW18-20 (n = 3 independent samples) and immunopanned for O4+ oligodendrocyte lineage cells, resulting in 2D cultures enriched for oligodendrocyte markers (*SOX10, OLIG2, PDGFRA*, and *MAG*) and depleted of astrocyte (*AQP4*) and neuron markers (*STMN2*) as compared to unpanned tissue (**Figures S4A– S4C**). We then treated cultures with vehicle or TIC for 14 days (**Figure 3G**). TIC treatment reduced the expression of MFOL/MOL marker MBP by approximately 15-fold in 2D culture as quantified by immunocytochemistry (mean ± SEM: vehicle = 0.091 ± 0.018, TIC = 0.0054 ± 0.002; **Figures 3H** and **3I**), confirming that organoid responses mirror those observed in primary human tissue.

### Acquisition of disease-associated signatures

We next investigated if the transcriptional shift we observed after TIC treatment might also be associated with the acquisition of disease signatures. Reactive^40,41^ or disease-associated oligodendrocytes (reported as DOL^42^, DAO^31^, immune OPCs/OLs^43,44^, and DA^45,46^) have been gaining attention, particularly in MS research. To understand what processes drive the transcriptional shift observed in TIC-treated oligodendrocyte lineage cells, we first performed Gene Ontology (GO) enrichment analysis. In TIC-treated oligodendrocytes, we observe an upregulation of pathways related to DNA damage response, proteasome function, regulation of mTORC1 signaling, and insulin responsiveness, alongside downregulation of oxidative phosphorylation, mRNA processing and translation, and cholesterol biosynthesis (**Figure 4B**; differentially expressed genes for each group listed in **Table S1J**, all significantly altered GO processes per group listed in **Table S1L**). Collectively, these changes are consistent with transcriptional programs previously reported in disease-associated oligodendrocytes and suggest a shift toward stress-responsive and metabolically compromised states. Indeed, more mature MFOL/MOL from TIC-treated assembloids showed decreased expression of the lactate transporter *MCT1* (*SLC16A1*), a key mediator of neuronal metabolic support that is reduced in patient tissue from neurological disorders including ALS^47^ and epilepsy^48^, alongside downregulation of genes associated with cholesterol synthesis pathways indicating disruption of lipid metabolism essential for oligodendrocyte health and myelination^26,27^ (**Figure 4C**).

**Figure 4.**
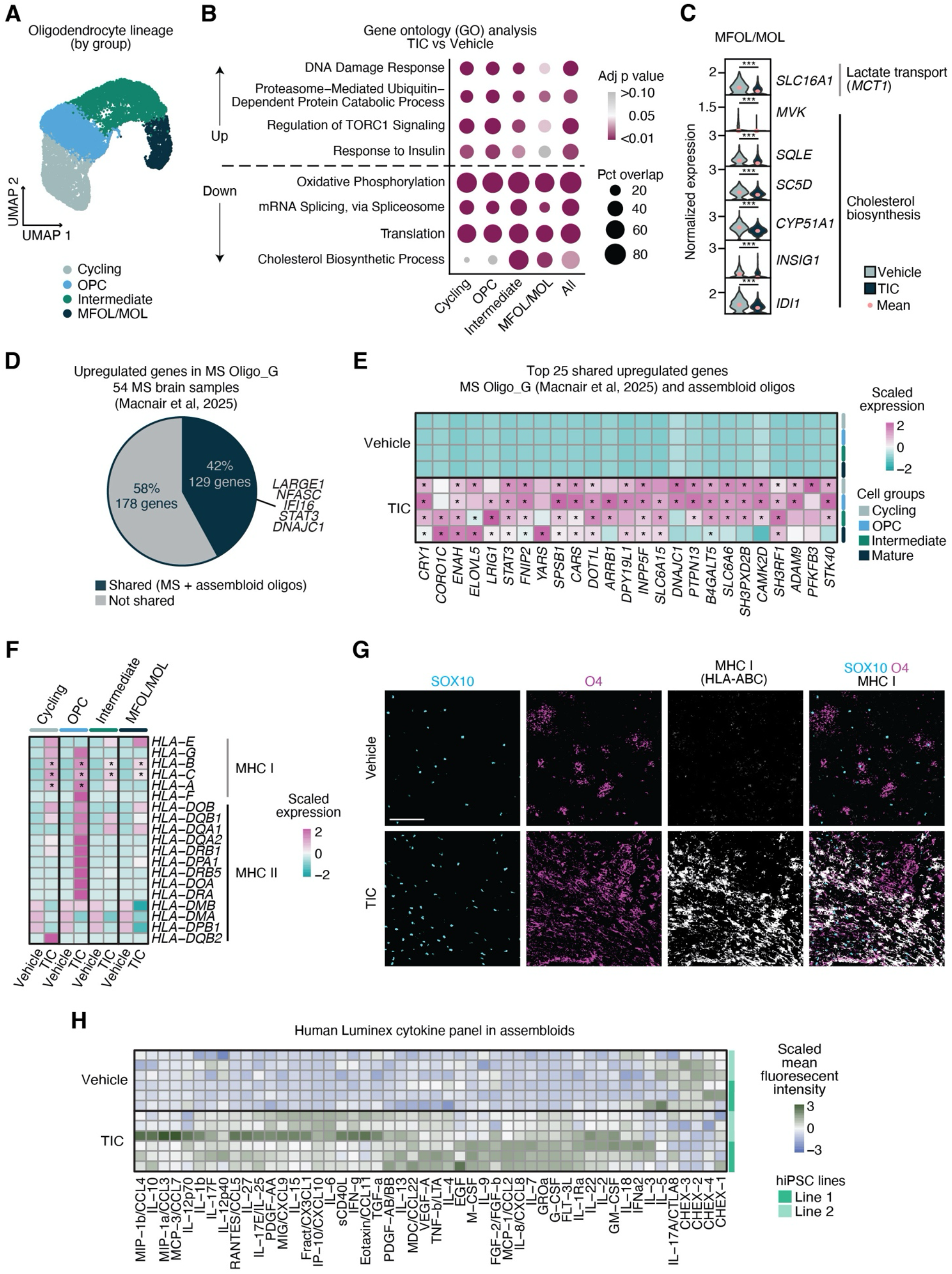
Disease-associated signatures in cytokine-treated hSpO oligodendrocyte lineage cells. **(A)** UMAP plot showing organoid-derived oligodendrocyte lineage cells treated with vehicle or TIC (n = 15,330 cells from 2 hiPSC lines). Seurat clusters were combined into groups for analysis. See **Figure 3C** for annotated Seurat clusters. **(B)** Gene ontology (GO) biological process analysis of genes upregulated or downregulated in TIC-treated oligodendrocyte lineage cells. Selected GO terms are shown for five groups: the four oligodendrocyte lineage subgroups and all oligodendrocyte lineage cells combined. Dot color indicates statistical significance (adjusted p value), and dot size represents the percentage of overlapping genes contributing to each GO term (0–100%). Terms were manually selected from the top significantly enriched categories for disease-signature relevance; full GO results are provided in **Table S1L**. **(C)** Violin plots showing *MCT1* lactate transporter and cholesterol biosynthesis marker (*SLC16A1, MVK, SQLE, SC5D, CYP51A1, INSIG1, IDI1*) gene expression in vehicle or TIC treated MFOL/MOL cells. Y-axis represents normalized gene expression, log transformed (counts per 10,000). Wilcoxon rank sum test; *SLC16A1* p <0.0001, *MVK* p = 0.0001, *SQLE* p <0.0001, *SC5D* p <0.0001, *CYP51A1* p <0.0001, *INSIG1* p = 0.0001, *IDI1* p <0.0001. Statistical significance is denoted as p < 0.001 (***). **(D)** Pie chart representing significantly upregulated genes in disease-associated MS oligodendrocyte cluster Oligo_G from MS patient dataset Macnair et al, 2025. Number/percent of genes in dark blue portion represents genes commonly upregulated in oligodendrocytes from hSpO/assembloids treated with TIC. Arrow shows example genes in common (*LARGE1, NFASC, IFI16, STAT3, DNAJC1*). All genes have adjusted p-value or false discovery rate (FDR) of less than 0.05 and greater than 0.25 log_2_ fold change over their respective controls. **(E)** Heatmap of top 25 markers from disease-associated MS oligodendrocyte cluster Oligo_G which are also upregulated in oligodendrocytes from TIC-treated assembloids. Cells within each group were pseudobulked by treatment. Gene expression was normalized to counts per 10,000, log-transformed, and then normalized to the average vehicle expression for each gene. Z-scores were computed within each gene across conditions. Wilcoxon rank sum test; (*) represents adjusted p-value of less than 0.05. **(F)** Heatmap showing MHC Class I and II gene expression across cell groups, split by condition. Cells within each group were pseudobulked by treatment. Gene expression was normalized to counts per 10,000, log-transformed, and then normalized to the average vehicle expression for each gene. Z-scores were computed within each gene across conditions. Wilcoxon rank sum test; (*) represents adjusted p-value of less than 0.05. **(G)** Representative image showing immunohistochemistry for MHC Class I (HLA-ABC) colocalized with SOX10 and O4 in sections of vehicle or TIC-treated assembloid. **(H)** Luminex-based multianalyte profiling of 48 cytokines and chemokines in supernatants from assembloid cultures after 8 weeks of vehicle or TIC treatment. Cytokine/chemokine levels are shown as log_2_ fold change of mean fluorescent intensity (MFI) relative to the average value of vehicle treatment per line. Values were Z-score-scaled on a per cytokine/chemokine basis, with green indicating high levels and purple indicating low values. n = 3 samples per line per condition and 2 assembloids per media sample in 2 hiPSC lines. Scale bar: 100 μm (**G**).

Examination of genes upregulated in TIC-treated oligodendrocyte lineage cells indicated that a subset of them was shared with genes upregulated in astrocytes, including *MTRNR2L12*, a stress-associated humanin-like transcript, the Wnt coreceptor *LRP6*, and the oligodendrocyte epigenetic regulator *CHD7*. On the other hand, some genes appeared to be selectively upregulated in oligodendrocytes, including the circadian transcription factor *NPAS2*, the growth factor *IGF2*, and the long noncoding RNA *H19* (**Figure S3L** and **Table S1K**). Further analysis of genes upregulated in the oligodendrocyte lineage upon TIC treatment revealed shared signatures with disease-associated oligodendrocytes previously reported in Macnair et al (2025), the largest cohort of MS brain samples to date^45^. We observed that 42% of the genes upregulated in the MS-specific oligodendrocyte cluster Oligo_G were also upregulated in our TIC-treated oligodendrocyte lineage cells. Shared genes included the heat shock protein *DNAJC1*, the demyelinating neuropathy-associated *NFASC*, and the gliosis regulator *STAT3* (**Figures 4D, E** and **S5A, B**), further supporting their reactive-like state.

A defining feature of disease-associated oligodendrocytes in MS and other CNS pathologies is the expression of Major Histocompatibility Complex (MHC) genes^10^. To determine whether our model system recapitulates this hallmark of immune-competent oligodendrocytes, we first examined MHC expression in TIC-treated hSpO-derived oligodendrocytes in our single-cell RNA sequencing dataset. TIC treatment induced robust expression particularly of MHC Class I genes across the oligodendrocyte lineage, with the highest expression observed in OPCs (**Figure 4F**).

We next confirmed MHC Class I expression at the protein level via immunohistochemistry, which also indicated extensive colocalization of MHC I with oligodendrocyte lineage cells expressing O4 (**Figure 4G**).

To determine whether this response to TIC treatment is biologically relevant, we assessed MHC induction in oligodendrocyte-enriched cultures immunopanned from human fetal cortical tissue (GW 18-20, n = 3 samples). Cultures were treated with vehicle or TIC for 14 days. Because previous work has shown the potent induction of MHC expression by IFN-γ in oligodendrocytes in rodent brain cultures^49^, purified rat oligodendrocyte cultures^50^, and autopsied human brain white matter cultures^51^, we included this treatment as a positive control (**Figure S6A**). Bulk RNA sequencing indeed indicated a robust increase in MHC Class I and II expression in both TIC and IFN-γ treatment (**Figure S6D**; differential expression data for each treatment listed in **Table S1M**). We again confirmed this expression at the protein level using immunocytochemistry (**Figures S6E** and **S6F**). The number of MHC Class I expressing oligodendrocytes increased with TIC and IFN-γ treatment by 4-fold and 10-fold, respectively (**Figure S6G**). Similarly, MHC Class II expression in oligodendrocytes increased 13-fold and 29-fold with TIC and IFN-γ treatment, respectively (**Figure S6H**).

Finally, we asked whether TIC exposure in assembloids was associated with broader inflammatory signaling beyond the direct TIC treatment. Using a Luminex cytokine release assay of assembloid supernatants, we found that despite the absence of microglia, TIC-treated assembloids released a diverse array of inflammatory mediators. These included previously reported immune and disease-associated cytokines IFN-γ, IL-6, and IL-1β, indicative of a sustained, inflammatory-like environment (**Figure 4H**). Collectively, these findings demonstrate that hSpO-derived oligodendrocytes recapitulate a core immunopathological feature of disease oligodendrocytes, and that assembloid systems support complex, pro-inflammatory environments permissive of in vitro disease-associated oligodendrocyte states.

## Discussion

Oligodendrocyte lineage cells are critical contributors to central nervous system development and disease, yet they remain significantly understudied in human systems compared to neurons. Here, we elucidate oligodendrocyte development, functional properties, and disease-relevant states in human organoids and assembloids. We show that these platforms support multiple stages of the oligodendrocyte lineage and enable interrogation of developmental and pathological processes that are historically challenging to study in human multi-cellular environments.

Many in vitro approaches to oligodendrocyte generation in 3D rely on specialized culture conditions optimized for producing mature, myelinating cells, particularly in cortical organoids^52–55^. Our system generates cells across the oligodendrocyte lineage, allowing cell type-resolved analyses of oligodendrocyte responses. Lineage progression was preserved across culture contexts, including within assembloids containing cortical and muscle tissues, indicating that oligodendrocytes maintain their intrinsic differentiation programs while providing opportunities to investigate their interactions with diverse, physiologically relevant cell types.

Using assembloids, we demonstrate oligodendrocyte migration into oligodendrocyte-naïve cortical tissue, capturing a key developmental behavior that is difficult to model in human systems. Although such migration does not reflect a native spinal-to-cortical trajectory in vivo, it provides a tractable platform to study oligodendrocyte colonization dynamics and neuron–glia interactions in three-dimensional tissue. Consistent with this, monosynaptic rabies tracing revealed direct connectivity between neurons and oligodendrocyte lineage cells in hSpO. This work represents the first in vitro investigation of neuron-oligodendrocyte synapses. While the functional significance of these contacts remains incompletely understood, their presence supports emerging models of activity-dependent regulation of oligodendrocyte lineage development^56,57^ and pathological contexts in which neuron–glia synaptic interactions are co-opted, such as neuron–glioma synapses^58–60^. These findings position hSpO as a platform for interrogating human neuron-glia communication.

Under cytokine-induced neuroinflammatory conditions, we observed robust responses across the oligodendrocyte lineage, paralleling reactive astrocyte responses within the same tissues. Oligodendrocytes exhibited transcriptional changes consistent with stress-responsive and disease-associated states, including loss of transcriptional programs for key homeostatic functions such as lactate transport and cholesterol biosynthesis. Notably, inflammatory exposure was associated with increased OPC proliferation accompanied by reduced representation of mature-like oligodendrocytes^61^, a pattern reported in multiple neurological disorders, including MS^9^, ALS^62^, and spinal cord injury^63^. This bias towards OPC expansion and away from more mature oligodendrocyte generation is consistent with changes observed in glial-enriched cultures derived from primary progressive MS patient hiPSC lines^64^, and has important implications in demyelinating diseases, where suppression of oligodendrocyte maturation could compromise endogenous remyelination. These findings highlight important mechanistic questions regarding differentiation blockade versus mature cell loss, with distinct implications for therapeutic intervention.

Oligodendrocytes also acquired immune-like features, including upregulation of MHC class I and II molecules, consistent with disease-associated oligodendrocyte states reported in human MS tissue and animal models. Although cytokine-treated assembloid oligodendrocytes do not fully recapitulate adult MS transcriptional signatures, the induction of disease-associated markers, a phenotype that has not been previously reported in a human model, highlights the utility of this system for modeling early, inflammation-driven pathological states.

It remains unclear whether inflammatory signals act directly on oligodendrocytes or indirectly through cell-to-cell interactions. Previous studies have shown that reactive astrocytes can be detrimental to oligodendrocyte survival and function^35,65^. Future work will be needed to disentangle the direct versus indirect effects of TIC on the oligodendrocyte lineage. This complexity is further compounded by evidence that inflammatory environments can alter lineage plasticity in astrocytes and OPCs, including reports of fate transitions between these lineages^66–68^. Additionally, the sources of additional cytokines detected in the culture supernatant remain undefined. These signals may further drive such disease-associated phenotypes, for example IL-6 and IL-1β are known to reduce the proliferative capacity of oligodendroglia^69,70^.

Together, this work establishes hSpO and assembloids as platforms for studying human oligodendrocyte lineage biology, neuron-glia interactions, and early disease-relevant responses. Notably, we demonstrate neuron-oligodendrocyte synaptic connectivity in an in vitro human system, introduce a viral strategy to interrogate these interactions, and show that inflammatory exposure induces oligodendrocyte transcriptional programs that overlap with disease-associated signatures identified in MS patient tissue, extending observations not previously captured in in vitro models. These systems bridge the gap between reductionist in vitro models and postmortem human tissue, enabling mechanistic studies of oligodendrocyte dysfunction and providing a foundation for therapeutic discovery in neuroinflammatory disorders.

## Limitations

Organoids lack the spatial organization, long-range patterning cues, and regional specialization present in vivo, which may influence oligodendrocyte maturation and function. The oligodendrocyte lineage in hSpO and assembloids also represents a predominantly developmental state, whereas many neurodegenerative reference datasets derive from adult postmortem tissue with chronic pathology. This developmental mismatch, together with technical differences between single-cell and single-nucleus sequencing approaches, likely contributes to the incomplete overlap observed between assembloid and patient-derived disease signatures. Future studies directly comparing human developmental and adult oligodendrocyte states, including across matched inflammatory contexts, will be essential for clarifying how early pathological programs evolve into chronic disease-associated phenotypes. Additional limitations relate to inflammatory modeling and cellular composition. Microglia, which are key regulators of neuroinflammation and oligodendrocyte biology in vivo, are absent from the current system. Incorporation of microglia and refined inflammatory paradigms in future assembloid models will enable more precise dissection of cell-type–specific contributions to oligodendrocyte pathology.

## Acknowledgements

We would like to thank members of the Andersen, Birey and Sloan labs and members of the Emory Brain Organoid Hub for useful discussions and support. We would especially like to thank Arvin Sarkissian, Fikri Birey, Nisha Raj, and Francisco Alvarez for their perspectives throughout the generation of this manuscript. We would also like to thank Standford Human Immune Monitoring Center (HIMC) for assistance with the Luminex cytokine-release panel. Rabies viral tracing work was supported by the GT3 Core Facility of the Salk Institute with funding from NIH-NCI CCSG: P30 CA014195 and funding from NIH Grant 1U24NS140479. This work was supported by a National Institute of Neurological Disorders and Stroke (NINDS) Fellowship (5F31NS135955-02) and training grant (T32NS096050) to T.P., National Institute of Mental Health (5R01MH125956-05) and NINDS (5R01NS123562-05) grants to S.A.S., and by grants from the University Research committee of Emory University (to J.A.), and the Congressionally Directed Medical Research Programs Multiple Sclerosis Research Program award sponsored by The Assistant Secretary of Defense for Health Affairs (HT94252410875; to J.A.).

## Author contributions

T.P., M.B., and R.Z. performed organoid assembly and maintenance, live imaging, cryosectioning, immunohistochemistry, confocal imaging, and quantitative analyses. S.S.W. prepared and maintained all myoblast cultures. T.P., M.B., and S.S.W. performed viral infections. T.P. and S.M. carried out single-cell dissociation and cell counting for single-cell RNA sequencing. T.P. prepared libraries for single-cell RNA sequencing samples. E.J.H. and S.A.S. supported experimental design of cytokine-treatment of assembloid and primary cultures and subsequent interpretation of these experiments. T.P. performed primary human tissue dissociation, oligodendrocyte enrichment, culture treatment, and sample collection. T.P., M.B., and S.S.W. performed immunocytochemistry and T.P. and S.S.W. performed imaging of primary cultures. T.P. and M.B. performed RNA isolation of all samples. S.S.W. conducted library preparation for bulk RNA sequencing experiments. T.P. performed processing and analysis of single-cell RNA sequencing, bulk RNA sequencing, and MS patient data with input and oversight from J.A. T.P. and M.B. collected culture supernatants for cytokine release assays. T.P., M.B., and J.A. designed experiments and interpreted the data. T.P. and J.A. conceived the project and wrote the manuscript. J.A. supervised the work.

## Lead Contact

Further information and requests for resources and reagents should be directed to and will be fulfilled by the Lead Contact, Jimena Andersen.

## Methods

### Data and code availability

Standard analysis pipelines for bulk and single cell RNA sequencing data were used (see details below). All code is available from the lead contact upon request.

### Protocol availability

The Emory University Brain Organoid Hub generated and maintained many of the hiPSC lines and organoid differentiations used for these experiments. Detailed protocols and demonstration videos on hiPSC thawing, maintenance, passaging, and organoid differentiation are available at www.brainorganoidhub.com/protocols.

### Cell lines, reagents, and material catalog numbers

**Table S1** details which hiPSC lines were used for each experiment, the number of differentiations from which data were generated, and the number of organoids or assembloids analyzed (**Table S1A**). This information is also provided in the corresponding figure legends. **Table S1** also includes catalog numbers for all culture reagents (**Table S1B**), antibodies (**Table S1C**), viruses (**Table S1D**), plasmids (**Table S1E**), and sequences for primers (**Table S1F**) used in this study.

### Variability terminology

We accounted for three levels of variability in this study, outlined below:

- Line-to-line variability: hiPSC lines derived from different individuals; each cell line is represented by different shapes in graphs unless otherwise noted.
- Experiment-to-experiment (differentiation) variability: a differentiation experiment represents a batch of hiPSCs differentiated in parallel; differentiations are noted in **Table S1A**.
- Organoid-to-organoid or assembloid-to-assembloid variability: values for individual organoids or assembloids are shown in graphs unless specified otherwise.

### Culture of hiPSCs

The hiPSC lines used in this study were validated using standard methods as previously described^28,71,72^. Three hiPSC lines derived from fibroblasts (8858-1 and 0524-1) and umbilical cord blood (C4-1) collected from healthy subjects were used for experiments (see **Table S1A** for details of hiPSC lines and donor sex for each line). All lines were genotyped by SNP array to confirm genomic integrity and regularly screened for mycoplasma. hiPSCs were cultured on recombinant human vitronectin with daily media changes of Essential 8 medium (E8) or double volume of Essential 8 Flex medium on weekends (as described in detail at www.brainorganoidhub.com/protocols). hiPSC colonies were passaged with ReLeSR upon reaching 80–90% confluency. **Table S1B** includes catalog numbers for all cell culture reagents.

### Generation of hSpO from hiPSCs

Human spinal organoids (hSpO) were formed following previously published protocols^15^. hiPSC colonies at 80–90% confluency were detached from culture plates using Accutase and formed into 3D organoids in AggreWell 800 (STEMCELL Technologies, 34815) plates, after which they were dislodged into 10 cm plates containing dorsomorphin (DM, 2.5 μM), SB431542 (SB, 10 μM), and Y-27/Rock Inhibitor (10μM) in E6+ medium (Essential 6 medium supplemented with penicillin/streptomycin 1×). On days 3 and 4, hSpO received DM (2.5 μM), SB (10 μM), and CHIR-99021 (CHIR, 3 μM) in E6+. From day 5 to day 9, hSpO received FGF2 (10 ng/mL), EGF (20 ng/mL), retinoic acid (RA, 0.1 μM), and CHIR (3 μM) in NBA+ medium (Neurobasal-A, B27 Minus Vitamin A 1×, GlutaMAX 1×, penicillin/streptomycin 1×) every 1–2 days. From day 10 to day 17, hSpO received FGF2 (10 ng/mL), EGF (20 ng/mL), RA (0.1 μM), CHIR (3 μM), and smoothened agonist (SAG, 0.1 μM) in NBA+ every other day. From day 18 to day 23, hSpO received N-2 supplement (1:100), BDNF (20 ng/mL), cAMP (50 μM), L-ascorbic acid (L-AA, 200 μM), IGF-1 (10 ng/mL), and DAPT (2.5 μM) in NBA+ every other day. From day 24 to day 43, hSpO received N-2 (1:100), BDNF (20 ng/mL), cAMP (50 μM), L-AA (200 μM), and IGF-1 (10 ng/mL) in NBA+ every 2–3 days. After day 43, hSpO received N-2 (1:100), BDNF (20 ng/mL), cAMP (50 μM), L-AA (200 μM), and IGF-1 (10 ng/mL) in NBA+ every 3–4 days.

### Generation of hCO from hiPSCs

Human cortical organoids (hCO) were formed following previously published protocols^73^. hiPSCs were cultured and aggregated as above. Following dislodge, hCO received daily ∼70% media changes in DM (2.5 μM) and SB (10 μM) in E6+ for 5 days. From day 6 to day 17, hCO received EGF (20 ng/mL) and FGF2 (20 ng/mL) in NBA+ daily, skipping weekends. From day 17 to day 24, hCO received EGF (20 ng/mL) and FGF2 (20 ng/mL) in NBA+ every 2–3 days. From day 25 to day 43, hCO were treated with BDNF (20 ng/mL) and NT-3 (20 ng/mL) in NBA+ every 2–3 days. After day 43, hCO received unsupplemented NBA+ every 3–4 days.

### Generation of hCO-hSpO assembloids

To generate hCO-hSpO assembloids for migration assays, hCO and hSpO were generated separately and later assembled by placing them in close proximity on top of cell culture inserts (0.4 mm pore size; Corning 353090) that were positioned in 6-well plates containing 1.5 mL of medium underneath the insert. The medium used for assembly was NBA+ (see above) supplemented with BDNF (20 ng/mL), NT-3 (20 ng/mL), L-AA (200 μM), and cAMP (50 μM). Three assembloids were maintained per insert. 1mL of medium underneath the insert was changed every 2–3 days. hCO and hSpO were assembled between day 90 and day 100 (both regions were the same age).

### Generation of 3D hSkM

Human Skeletal Myoblasts (hSkM) from Thermo Fisher Scientific (Cat# A11440, Lot# 3253421) were maintained with Skeletal Muscle Cell Growth Medium supplemented with 50 μg/ml fetuin, 10 μg/ml Insulin, 0.4 μg/ml Dexamethasone, and 5% Fetal bovine serum in 10cm plates. Media was changed every other day, and cells were passaged using 0.25% Trypsin-EDTA when they reached 80% confluency. For 3D hSkM cultures, hSkM were dissociated using Trypsin and resuspended in Geltrex at a density of 2,400 hSkM per μl. 10 μl of cell suspension was aliquoted into silicone wells (Ibidi; Thermo Fisher Scientific 50-305-782) in 6-well tissue culture plates and incubated for 20-30 minutes at 37°C to allow Geltrex to gel up, then 4ml of Skeletal Muscle Cell Growth Medium was added. The next day, silicone wells containing hSkM were moved into new 6-well tissue culture plates coated with anti-adherence rinsing solution and the medium was changed every 2–3 days. After 7–10 days, media was changed to Skeletal Muscle Cell Differentiation Medium supplemented with penicillin/streptomycin 1× and 10 μg/ml insulin, with media changes every 2–3 days.

### Generation of hCO-hSpO-hSkM assembloids

To generate hCO-hSpO-hSkM assembloids, hCO, hSpO, and hSkM were generated separately and later assembled by placing them in close proximity on a cell culture insert as previously described. Two assembloids were formed per insert. For each assembloid, two hSpO were placed between one hCO and one hSkM to prevent direct contact between hCO and hSkM. The medium used for assembly was HS+ media (DMEM/F12 supplemented with 1% non-essential amino acids, 1% insulin-transferrin-selenium, 1% penicillin/streptomycin, L-ascorbic acid at 200 μM, and cAMP at 50 μM). Medium was changed every 2–3 days. At the time of assembly, hCO were day 90–120, hSpO were day 40–50, and hSkM were day 10–15 following differentiation media.

### Vehicle and TIC treatment of assembloids

TIC treatment consisted of HS+ media with TNF-α (30 ng/mL), IL-1α (3 ng/mL), and C1q (400 ng/mL) from freshly thawed aliquots added to media directly before media change. Vehicle-treated controls received HS+ medium supplemented with an equivalent volume of sterile 0.1% BSA in PBS, matching the cytokine solution. Assembloids were fused for 2 weeks before receiving 8 weeks of vehicle or TIC treatment.

### Human Tissue

All human tissue samples were obtained in compliance with policies outlined by the Emory School of Medicine IRB office.

### Single cell dissociation

We used a previously established protocol to create single cell suspensions from human organoids, assembloids, and primary human fetal cortical tissue (GW18-20)^74,75^. Briefly, we mechanically (razor blade) and enzymatically dissociated tissue using papain (Worthington Biochemical LS003126) at 20 U/mL at 37°C for 45 minutes to 1 hour. Organoids and assembloids were dissociated in batches of 2–5 per well in six-well plates. Papain was quenched with ovomucoid solution (Worthington Biochemical LS003086), and samples were triturated using P1000 and P200 pipettes to obtain single-cell suspensions.

### Immunopanning

To isolate cell types of interest, single-cell suspensions were added to a series of 10 cm (organoid and assembloid) or 15 cm (primary tissue) petri dishes pre-coated with cell type-specific primary antibodies for depletion of unwanted cell types and final enrichment of oligodendrocyte lineage cells and astrocytes (see **Figure S4A** for immunopanning scheme and **Table S1C** for all antibody details). Cell suspensions were incubated on plates for 5–25 minutes depending on the coating antibody, and unbound cells were then transferred to the next plate in the sequence. Positive-selection plates with attached cells were washed 5–7 times with PBS without calcium or magnesium to remove unbound cells. Plates were then incubated in trypsin solution (Sigma-Aldrich, T9935-100MG) at 37°C for 5 minutes, quenched with 30% FBS (Thermo Scientific 10082147) in DMEM, and cells were dislodged from the plates for 2D plating and culturing.

Cell solutions were pelleted at 200g for 5 minutes, then resuspended depending on downstream need. For culturing, cells were resuspended and counted with a hemocytometer. Cells were seeded at a density of 40,000 cells per cm^2^ in wells coated with poly-D-lysine and laminin (25 μg/mL) and fed with proliferation media (PDGF-AA at 10 ng/mL and FGF2 at 10 ng/mL in NBA+). A complete media change was performed 24 hours after plating, then half changes were performed every 2–3 days.

After one week, cells were fed with a full change of maturation media containing T3 (40 ng/mL) and ITS 1× in NBA+. TIC treatment consisted of maturation media with TNF-α (30 ng/mL), IL-1α (3 ng/mL), and C1q (400 ng/mL) added to media directly before media change. Vehicle-treated controls received maturation media supplemented with an equivalent volume of sterile 0.1% BSA in PBS, matching the cytokine solution. IFN-γ treatment consisted of maturation media with IFN-γ (0.06 μg/mL) added to media directly before media change.

### Organoid viral labeling and monosynaptic rabies tracing

See **Table S1D** for catalog number and concentration of viruses used: LV-SOX10MCS5-GFP, LV-SOX10MCS5-Cre, AAVDJ-CAG-FLEX-oG-WPRE-SV40pA, AAVDJ-CAG-FLEX-TVA-mCherry, and EnvA G-Deleted Rabies-eGFP. Viral infection was performed in batches of 3–6 organoids per well in a six-well plate. All media was removed from the well, and virus was diluted in hSpO media. 3 μL of viral solution per organoid was added to each well and incubated for 30 minutes at 37°C with plates tilted to ensure solution remained in contact with organoids. After 30 minutes, 200 μL hSpO media per organoid was added to each well, and after overnight incubation, an additional 200 μL hSpO media per organoid was added to the well, with plates remaining tilted in the incubator. After this, culture returned to regular organoid maintenance with media changes every 3–4 days. Rabies virus was added in the above-described manner after 2 weeks. AAVs were acquired from Addgene. Rabies viruses were obtained from the Salk Institute Viral Vector Core. Lentiviruses were acquired and generated with Addgene and VectorBuilder.

### Organoid cryopreservation and immunohistochemistry

Cryopreservation and immunohistochemistry in hSpO and hCO-hSpO-hSkM were performed as previously described with some modifications^15,72,75^. Briefly, organoids or assembloids were fixed in 4% paraformaldehyde (PFA in PBS, Electron Microscopy Sciences 15710) for 1 hour. Fixation was followed by three PBS washes. Organoids and assembloids to be cryosectioned were placed in 30% sucrose solution in PBS for 24–48 hours or until samples sank to the bottom, embedded in 1:1 30% sucrose:OCT solution (Tissue-Tek OCT Compound 4583) in molds (Fisher 22-363553), and frozen at -20°C. For immunohistochemistry, 15–20 μm thick sections were cut using a cryostat (Leica) onto slides (VWR 48311-703). PAP pen (Fisher Scientific NC0822416) was used to create hydrophobic regions around cryosections. Cryosections were then washed with PBS to remove excess OCT, blocked for 1 hour at room temperature (10% normal donkey serum (Sigma S30-100ML), 0.3% Triton X-100 diluted in PBS), and incubated overnight at 4°C with primary antibodies in blocking solution without Triton (**Table S1C** lists all antibodies used in this study). The next day, cryosections were washed with PBS and then incubated with secondary antibodies in blocking solution without Triton for 1 hour at room temperature protected from light. Following washes with PBS, nuclei were visualized with Hoechst 33258 (1:7,000 applied for 5 minutes, Fisher Scientific H3569). Finally, slides were mounted for microscopy with coverglasses (Fisher Scientific S24461) using Aquamount solution (Polysciences 18606-20).

### Whole organoid immunohistochemistry

Whole organoids were fixed in 4% PFA in PBS for 1 hour at 4°C. Organoids were placed in blocking solution as described above for 4 hours, followed by primary antibody solution (blocking solution without Triton) for 2–3 days. Organoids were then washed three times for 5–10 minutes per wash, then placed in secondary antibody solution (blocking solution without Triton) overnight protected from light. Organoids were placed in Hoechst solution as described above for 5 minutes, then washed three times for 15 minutes per wash before placed on slides with coverglasses (Fisher Scientific S24461) using Aquamount (Polysciences 18606-20) in 2-4 stacked iSpacers (SunJin Lab IS213) per slide.

### Imaging

Slides, 2D culture plates, and live cultures were imaged on a Leica Stellaris 5 confocal microscope. Live imaging was performed under environmentally controlled conditions (37°C, 5% CO2) in the same microscope.

### Human primary fetal 2D immunocytochemistry

Glia-enriched 2D cells were fixed in 4% PFA for 10 minutes at 4°C. Cells were then washed three times in PBS and incubated for 10 minutes in blocking solution as described above, then incubated overnight at 4°C with primary antibodies in blocking solution without Triton. The next day, cells were washed with PBS and then incubated with secondary antibodies in blocking solution without Triton for 30 minutes at room temperature. Following washes with PBS, nuclei were visualized with 5 minutes incubation of Hoechst solution as described above. Cells were imaged in the culture plate in PBS.

### Migration quantification

Quantification of migrated cell area was carried out using the image analysis software Fiji^76^. Area of migrated cells (GFP in hCO) was normalized to the starting area of GFP cells in hSpO at day 2, and the same assembloids were imaged overtime. For each image, first the Despeckle feature was used, and background was removed by adjusting the minimum threshold to remove the lowest intensity pixels (same adjustment for all images within a given experiment). A border was drawn around the hSpO at day 2, and a mask was created of the GFP channel within that border. Then, a border was drawn around the hCO at days 2, 7, and 14, and background signal from the culture insert outside of this border was removed. For each day, a mask was created of the GFP channel, and area of GFP in hCO was measured and normalized to day 2 starting GFP area in hSpO.

### Immunohistochemistry and immunocytochemistry quantification

MBP area quantification for 2D cells was carried out using the image analysis software Fiji. First, maximum projection from all z-stacks was generated. The Despeckle feature was used, then background was removed by adjusting the minimum intensity to remove the lowest intensity pixels (same adjustment for all images within a given experiment). A mask was created of the MBP or Hoechst channel to restrict our measurement to the marker of interest, with the same threshold used for each marker within a given experiment.

Cell counting in rabies tracing and MHC experiments was performed with QuPath^77^. First, maximum projection from all z-stacks was generated. The Despeckle feature was used, then background was removed by adjusting the minimum threshold to remove the lowest intensity pixels (same adjustment for all images within a given experiment). Total GFP+ or SOX10+ cells with a Hoechst+ nucleus were counted to obtain the total population, then the colocalized marker was overlaid, and the number of double-positive cells were counted (O4, MAP2, or mCherry depending on the experiment).

### RNA collection, isolation, and Real-time quantitative PCR

For bulk RNA-seq of hSpO, organoids were placed in 1.7mL tubes, all media was removed, then samples were placed on dry ice and stored at - 80°C until RNA isolation. For bulk-RNAseq of 2D primary samples, 500 μL Qiazol (Qiagen 79306) was added to each well, pipetted 10-20 times, then moved to 1.7 mL tube and snap frozen on dry ice. Samples were stored at -80°C until RNA isolation. mRNA was isolated using the RNeasy Mini Kit and RNase-Free DNase Set (QIAGEN 74106) and stored at -80°C. Template cDNA was prepared by reverse transcription using the Maxima First Strand cDNA Synthesis Kit (Life Technologies K1672) and stored at -20°C. qPCR was performed using SYBR Green (Life Technologies 4312704) on a QuantStudio Real-Time PCR Flex System. All primers used are listed in **Table S1F**.

### SOX10-MCS5-GFP and SOX10-MCS5-Cre virus generation

Plasmid LV-SOX10MCS5-GFP was obtained from Addgene (Plasmid# 115783) as a bacterial agar stab. The bacteria containing the plasmid was grown on 10 cm LB agar (US Biological, L1515) plates containing 0.1mg/ml ampicillin (Sigma, A9518) overnight at 37°C. The next day, a single bacterial colony was picked from the agar plate using a sterile bacterial loop and grown in 50 ml LB broth (Thermo Fischer Scientific, BP9731-500) containing 0.1 mg/ml ampicillin. Bacteria were grown for 17–18 hours at 37°C on a shaker maintained at 250 rpm. The plasmid DNA was purified using a Miniprep kit (Qiagen, 27104). The concentration of eluted DNA was measured using Qubit Fluorometric Quantification. The DNA was normalized to 30 ng/μl to a total volume of 11 μl in elution buffer (Qiagen, 27104) and sent to plasmidsaurus for sequencing. Once the sequence of the plasmid DNA was confirmed, lentivirus packaging was performed by VectorBuilder. The SOX10-Cre plasmid was generated by replacing the GFP sequence from Addgene plasmid# 115783 with a Cre-recombinase sequence from VB241003-1053spr by VectorBuilder. This lentivirus was then packaged by VectorBuilder’s mini-scale packaging service from the generated plasmid.

### Cytokine-release luminex immunoassay

At 8 weeks of cytokine treatment (hSpO d120, same timepoint as single cell), 1 mL of assembloid supernatant was removed from each well (2 assembloids per well on insert) for three wells per condition (vehicle or TIC) in two hiPSC lines. Samples were snap-frozen on dry ice, then shipped to the Stanford Human Immune Monitoring Center (HIMC). Samples underwent Luminex 48-plex cytokine detection panel (H80-Panel 1, HCYTA-60K-PX48). Raw values were returned as mean fluorescent intensity (MFI) and processed according to HIMC recommendation. All sample values were normalized to the average vehicle MFI value for that hiPSC line. Heatmaps are presented as z-scores computed within each cytokine.

### Single cell RNA sequencing analysis and quality control

All cultures for single cell datasets generated in this study were maintained, isolated, and processed concurrently. hSpO and assembloids were dissociated into single cell solution as described above. Dissociated cells were resuspended in ice-cold PBS containing 0.02% BSA and loaded onto a GEM-X 3’ chip to generate gel beads in emulsion (GEMs). Single-cell RNA sequencing libraries were prepared with Chromium GEM-X Single Cell 3’ Reagent Kits v4 (PN-1000691) according to manufacturer instructions. Libraries from different samples were pooled and sequenced by Admera Health on a NovaSeq X Plus 25B (Illumina) using 150 × 2 chemistry, targeting 50,000 reads per cell. Demultiplexing, alignment, barcode and UMI counting, and aggregation were performed using Cell Ranger (cellranger v8.0.1 with default settings), which provided a feature-by-count matrix. Further analysis was performed using the R package Seurat (v5.0) from the Satija Lab^78^. We first excluded cells with more than 9,000 or fewer than 1,000 detected genes, as well as those with mitochondrial content higher than 8%. Gene expression was then normalized using the default global-scaling normalization method (normalization.method = “LogNormalize,” scale.factor = 10,000, selection.method = “vst” in Seurat) and scaled (mean = 0 and variance = 1 for each gene, as recommended by Seurat) prior to principal component analysis (PCA). The top 17 principal components were utilized for clustering with a resolution of 0.15, implemented using the FindNeighbors() and FindClusters() functions in Seurat.

We performed doublet removal using scDblFinder() to score all cells and remove cells with the highest doublet scores at a rate of 0.4% per 1,000 cells in each sample, the doublet rate predicted by 10x Genomics. We then performed more stringent quality control, filtering out cells with fewer than 1,300 unique genes and more than 7% mitochondrial gene expression. After doublet and low-quality cell removal, we again performed normalization and log transformation on raw counts as described above. Due to sex differences between hiPSC lines, we integrated by cell line using the Harmony() function. The top 17 principal components were utilized for clustering with a resolution of 0.35, implemented using the FindNeighbors() and FindClusters() functions in Seurat. We identified subclusters of neurons, oligodendrocytes, astrocytes, and fibroblasts based on expression of known markers (eg. *STMN2, OLIG2, HES1, COL1A1*, respectively).

After labeling all cells into subclusters, **Figure 1** oligodendrocyte lineage UMAP was generated using hSpO and vehicle-treated assembloid samples in the oligodendrocyte subcluster. **Figure 3** oligodendrocyte lineage UMAP was generated using vehicle-treated and TIC-treated assembloid samples in the oligodendrocyte subcluster. **Figure S3** astrocyte UMAP was generated using vehicle-treated and TIC-treated assembloid samples in the astroglia subcluster. In each UMAP, stressed clusters expressing *VEGFA, DDIT3, XBP1*^79^, or more than 1.25% ribosomal gene expression were removed. Oligodendrocyte subclusters were processed with 17 PCAs at a resolution of 0.35. The astroglia subcluster was processed with 17 PCAs at a resolution of 0.2.

#### Differential expression

Differentially expressed genes were determined using the Seurat function FindMarkers() with the Wilcoxon rank-sum test.

#### Label transfer from primary reference

To perform label transfer from the Andersen, Thom et al, dataset^19^, we excluded nuclei and utilized cells-only data from the oligodendrocyte lineage UMAP. We used Seurat’s “Mapping and annotating query datasets” vignette. After finding anchors, we used the TransferData() function to classify query cells based on human spinal cord cell type labels. TransferData() returns a matrix with predicted IDs and prediction scores, which we added to the organoid and assembloid oligodendrocyte metadata.

#### Reactive astrocyte signature

Seurat AddModuleScore() was used to calculate the average expression levels of each module signature on a single-cell level, subtracted by the aggregated expression of control feature sets. All analyzed features are binned based on averaged expression, and control features are randomly selected from each bin. Genes for the reactive astrocyte module came from Liddelow et al, 2017^35^ **Figure 1A** “Pan reactive” markers *LCN2, STEAP4, S1PR3, TIMP1, HSPB1, CXCL10, CD44, OSMR, CP, SERPINA3, ASPG, VIM*, and *GFAP*^35^.

#### Gene ontology (GO)

Differentially expressed genes were uploaded to Enrichr^80^, and “GO Biological Process 2025” data were exported.

#### Pseudobulk aggregation

To generate heatmaps of average gene expression per cell group and condition, we used the Seurat function AggregateExpression() to calculate pseudobulk counts for each grouped cluster and treatment combination. Expression values of 0 were replaced with 1e-12 so that all values could be normalized to the vehicle expression of that cluster for each gene.

#### Disease dataset comparison

Markers for disease-associated oligodendrocyte clusters (Oligo_F and Oligo_G) were obtained from Macnair et al, 2025^45^. For Venn diagram analyses, genes from the Macnair dataset were included if they met significance thresholds of false discovery rate (FDR) < 0.05 and absolute log_2_ fold change (log_2_FC) ≥ 0.25. TIC-induced differentially expressed genes in assembloid oligodendrocytes were defined using an adjusted p-value < 0.05 and absolute log_2_FC ≥ 0.25.

For heatmap visualization of shared upregulated genes, the top 25 marker genes for the Macnair cluster Oligo_G that were also significantly upregulated in cytokine-treated assembloids (adjusted p-value < 0.05 and log_2_FC ≥ 0.25) were selected and plotted.

### Bulk RNAseq library preparation and analysis

Concentration of the eluted RNA was measured using Qubit Fluorometric Quantification. The library was prepared using Magic prep NGS (Tecan 30186621) according to manufacturer recommendations with 150 ng RNA. The prepared libraries were sent to Admera for sequencing (NovaSeq X Plus 10B) targeting approximately 40 million paired end reads per sample.

FASTQ files were trimmed using Trimmomatic, aligned to hg19 with STAR, and reads were summarized with featureCounts. Raw read counts were converted to log_2_ counts per million (log_2_CPM). Bulk RNA-sequencing differential expression analysis was performed using the limma package with the voom transformation in R. The mean–variance relationship was modeled using voom, generating precision weights for each observation. Weighted linear models were then fit for each gene using limma, followed by specification of contrasts of interest via a contrast matrix. Empirical Bayes moderation was applied to the linear model fits to improve variance estimation across genes. Differentially expressed genes (DEGs) were identified from the moderated statistics using the topTable function, with p-values adjusted for multiple testing using the Benjamini– Hochberg false discovery rate (FDR) method. P-values were then doubled as a further Bonferroni adjustment for multiple comparison (TIC vs Vehicle, IFN-γ vs Vehicle).

### Schematic preparation

Schematics were prepared with Adobe Illustrator and Biorender.com.

### Statistical analysis

GraphPad Prism was used for statistical analysis of all immunohistochemistry and immunocytochemistry quantification. All data underwent normality testing prior to selecting appropriate statistical tests. Figure legends detail exact statistical tests used for each dataset. Data represent mean ± SEM unless otherwise noted. Statistical significance is denoted as p < 0.05 (*), p < 0.01 (**), and p < 0.001 (***) unless otherwise noted.

## Supplemental Information

**Figure S1.**
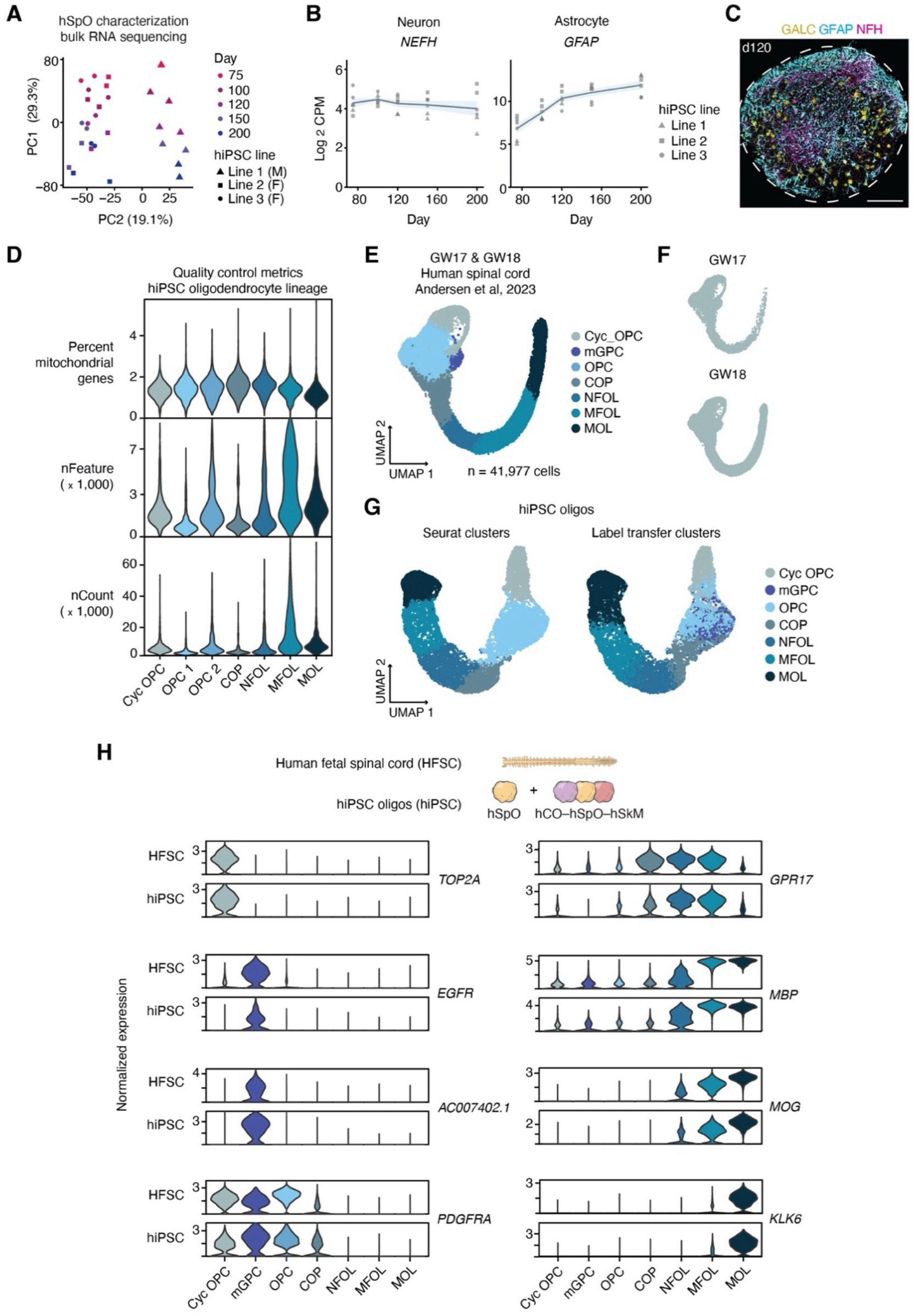
Validation of oligodendrocyte lineage identity and correspondence with primary human fetal spinal cord. **(A)** PCA plot showing PC1 and PC2 in bulk RNA sequencing hSpO samples over time (days 75 to 200). n = 2 replicates shown per hiPSC line and timepoint, with 3 hiPSC lines and 4 timepoints. Each replicate consists of 2-3 pooled organoids. **(B)** Log_2_ Counts Per Million (CPM) expression of canonical neuron (*NEFH*) and astrocyte (*GFAP*) markers from day 75 to day 200. n = 2 replicates shown per hiPSC line and timepoint, with 3 hiPSC lines and 4 timepoints. Each replicate consists of 2-3 pooled organoids. **(C)** Representative immunohistochemistry showing oligodendrocytes (GALC), astrocytes (GFAP), and neurons (NFH) in day 120 hSpO whole organoid. **(D)** Violin plots of quality control metrics showing percent of mitochondrial genes, number of features (nFeature, genes), and number of UMI counts (nCount) per cell in each oligodendrocyte lineage cell type. **(E)** UMAP visualization of the primary human spinal cord oligodendrocyte lineage at Gestational Weeks (GW) 17 and 18 (n = 41,977 cells from Andersen, Thom et al, 2023). Cells colored by cell type. **(F)** UMAP plot showing oligodendrocyte lineage from primary human spinal cord separated by age. **(G)** UMAP plots showing a comparison of organoid-derived oligodendrocyte lineage cells colored by Seurat annotated cluster (left UMAP) or by label transfer annotated clusters (right UMAP; label transfer from Andersen, Thom et al, 2023). **(H)** Violin plots showing gene expression of selected oligodendrocyte lineage marker genes compared between primary human fetal spinal cord (HFSC) clusters and organoid-derived (hiPSC) Seurat annotated clusters. Y-axis represents normalized gene expression, log transformed (counts per 10,000). Scale bar: 200 μm (**C**).

**Figure S2.**
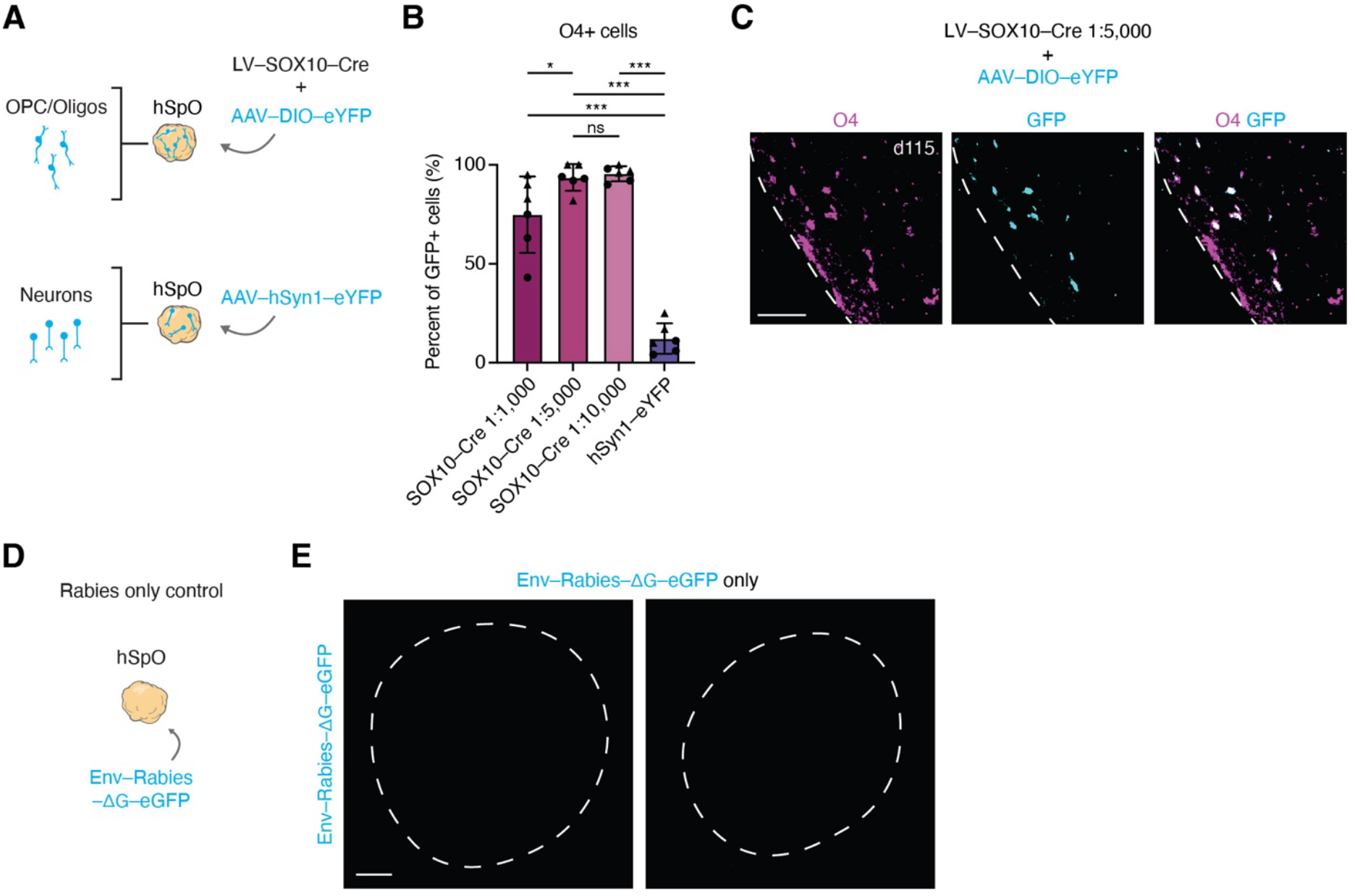
Validation of monosynaptic rabies tracing approach in hSpO. **(A)** Schematic detailing hSpO infection with either LV-SOX10-Cre + AAV1-EF1a-DIO-eYFP to label oligodendrocytes or AAV-PHP.eB-hSyn1-eYFP to label neurons as a negative control. **(B)** Quantification of O4+ cells colocalizing with eYFP in hSpO infected with either LV-SOX10-Cre + AAV-DIO-eYFP or AAV-hSyn1-eYFP. LV-SOX10-Cre was used at concentrations of 1:1,000, 1:5,000 or 1:10,000. Mean ± SEM: [1:1,000] = 74.8% ± 7.9%, [1:5,000] = 93.5% ± 2.7%, [1:10,000] = 95.5% ± 1.5%, [hSyn1-eYFP] = 12.2% ± 3.2%; n = 6 organoids from 2 hiPSC lines, with 2 sections quantified and averaged per organoid; one-way ANOVA with Tukey’s multiple comparisons: 1:1,000 vs 1:5,000 p = 0.039, 1:1,000 vs 1:10,000 p = 0.020, 1:1,000 vs hSyn1–eYFP p < 0.0001, 1:5,000 vs 1:10,000 p = 0.989, 1:5,000 vs hSyn1-eYFP p < 0.0001, 1:10,000 vs hSyn1-eYFP p < 0.0001). Statistical significance is denoted as p < 0.05 (*), p < 0.01 (**), and p < 0.001 (***). **(C)** Representative immunohistochemistry image of GFP and O4 colocalization in LV-SOX10-Cre (1:5,000) and AAV-DIO-eYFP. **(D)** Schematic detailing hSpO infection with only enveloped G-deleted rabies-eGFP (Env-Rabies-ΔG-GFP) as a negative control. **(E)** Representative images of two hSpO infected with only enveloped G-deleted rabies-eGFP (Env-Rabies-ΔG-GFP). Scale bar: 50 μm (**C**), 200 μm (**E**)

**Figure S3.**
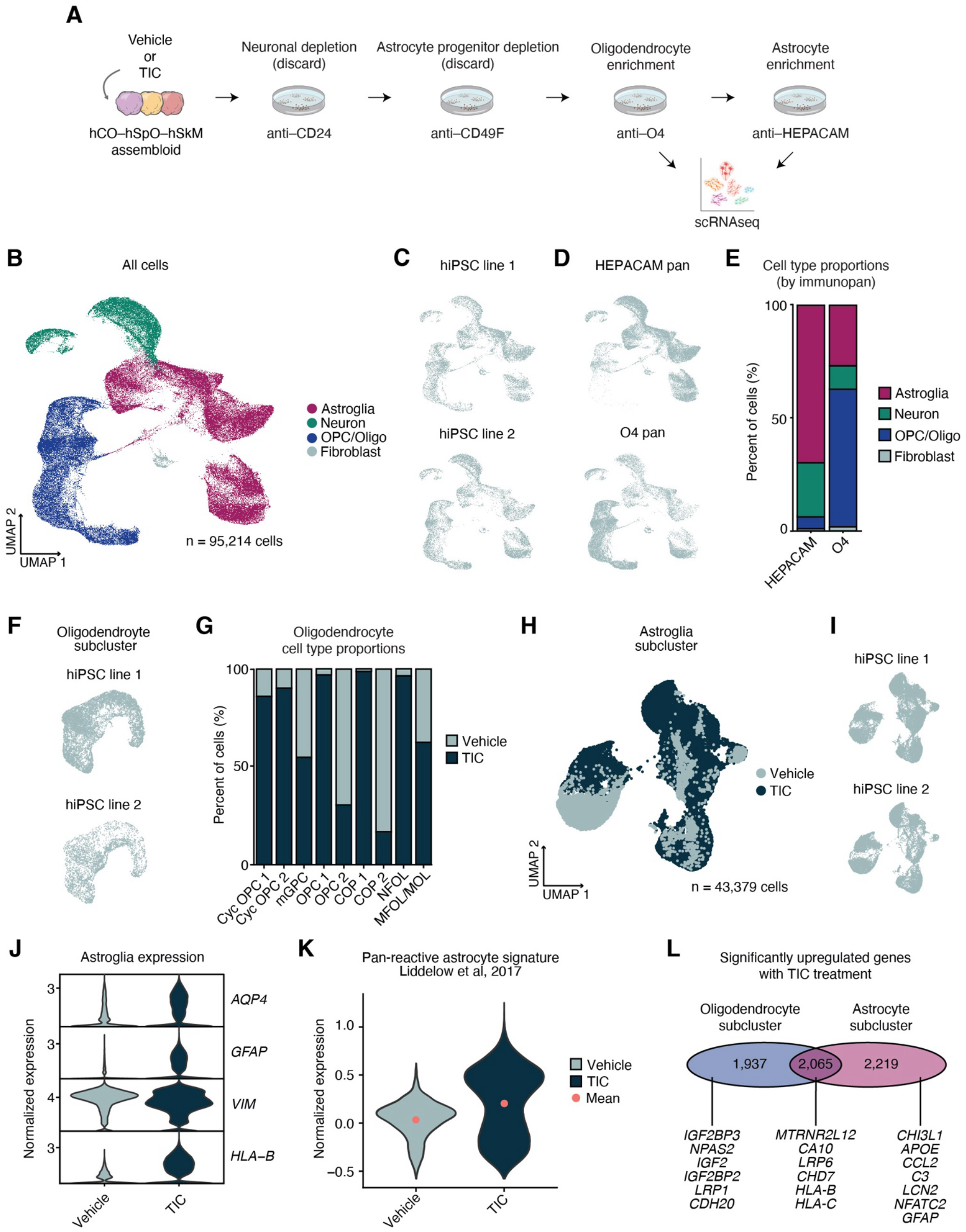
Astrocyte and oligodendrocyte lineage immunopanning and single cell RNA sequencing validation. **(A)** Schematic detailing assembloid culture treated with vehicle or TNF–α, IL–1α, and C1q (TIC) for 8 weeks (hSpO day 120, hCO–hSpO–hSkM 10 weeks of assembly). Immunopanning scheme depleted CD24+ cells (neurons), CD49F+ cells (astrocyte progenitors), then enriched for O4+ cells (oligodendrocytes) or HEPACAM+ cells (astrocytes). Cells in O4 and HEPACAM-enriched plates were separately captured for single cell RNA sequencing (scRNAseq, 10x Chromium). **(B)** UMAP plot showing organoid-derived cells following O4 and HEPCAM immunopanning in vehicle and TIC-treated groups (n = 95,214 cells from 2 hiPSC lines) colored by cell type. **(C)** UMAP plot of all cells separated by the hiPSC line they were derived from. **(D)** UMAP plot of all cells separated by the immunopan isolation method they were derived from. **(E)** Stacked bar plot showing the cell type enrichment as a percentage of all captured cells in each of the immunopan isolation methods. **(F)** UMAP plot showing subclustered oligodendrocyte lineage cells separated by the hiPSC line they were derived from. See **Figures 3B** and **3C** for clustering by treatment condition or cell type. **(G)** Stacked bar plot showing the percentage of cells in the vehicle or TIC treatment groups in each oligodendrocyte lineage cluster. **(H)** UMAP plot showing subclustered organoid-derived astroglia cells grouped by treatment (n = 43,379 cells from 2 hiPSC lines). **(I)** UMAP plot showing astroglia separated by the hiPSC line they were derived from. **(J)** Violin plot showing marker genes for astroglia and astrocyte reactivity in vehicle and TIC-treated astroglia. Y-axis represents normalized gene expression, log transformed (counts per 10,000). **(K)** Violin plot of pan-reactive astrocyte signature genes upregulated in astroglia after TIC treatment. Pan-reactive signature genes were identified from Liddelow et al, 2017 and added to organoid-derived dataset using the AddModuleScore() function in Seurat. **(L)** Venn diagram comparing number of differentially expressed genes following TIC treatment between oligodendrocyte (left) and astrocyte (right) subclusters. Center number in the overlap represents number of shared genes between oligodendrocytes and astrocytes. Arrows indicate representative genes uniquely altered in oligodendrocytes (left), uniquely altered in astrocytes (right), or altered in both lineages (center).

**Figure S4.**
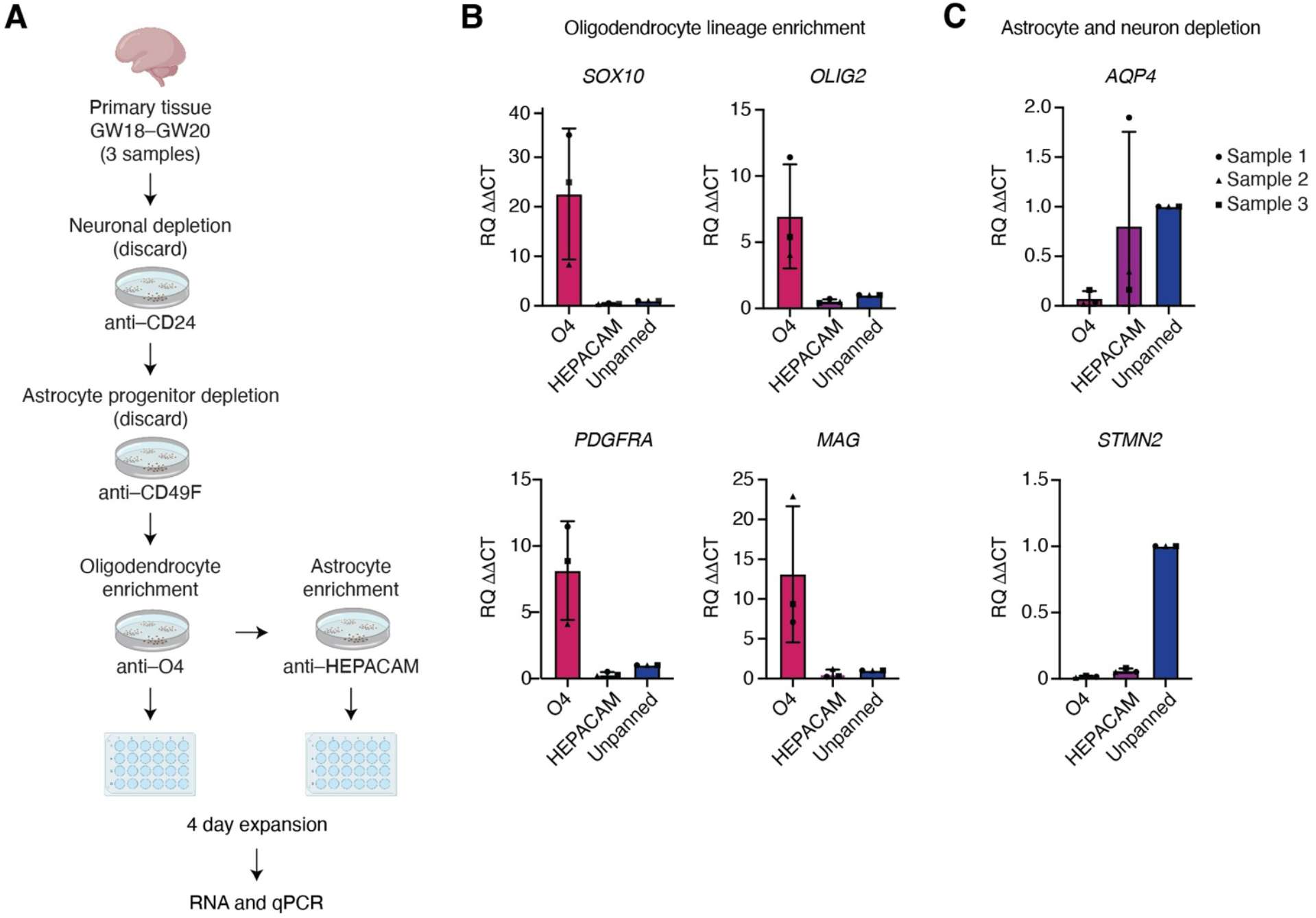
Validation of immunopanning isolation method in primary tissue. **(A)** Schematic detailing oligodendrocyte enrichment from human primary fetal tissue. Immunopanning scheme depleted CD24+ cells (neurons), CD49F+ cells (astrocyte progenitors), then enriched for O4+ cells (oligodendrocytes) or HEPACAM+ cells (astrocytes). Cells in O4 and HEPACAM-enriched plates were separately cultured for 4 days, then underwent RNA isolation and quantitative PCR (qPCR). **(B)** qPCR showing expression of oligodendrocyte-specific genes following immunopanning. Data is shown as ΔΔCt normalized to housekeeping gene *ACTIN* (ΔCt) then normalized to unpanned tissue sample of the same gestation week. n = 3 samples GW18–GW20, 1 replicate per immunopan condition. **(C)** qPCR showing expression of astrocyte (*AQP4*) and neuron (*STMN2*) marker genes following immunopanning. Data is shown as ΔΔCt normalized to housekeeping gene *ACTIN* (ΔCt) then to unpanned tissue sample of the same gestation week. n = 3 samples GW18–GW20, 1 replicate per immunopan condition.

**Figure S5.**
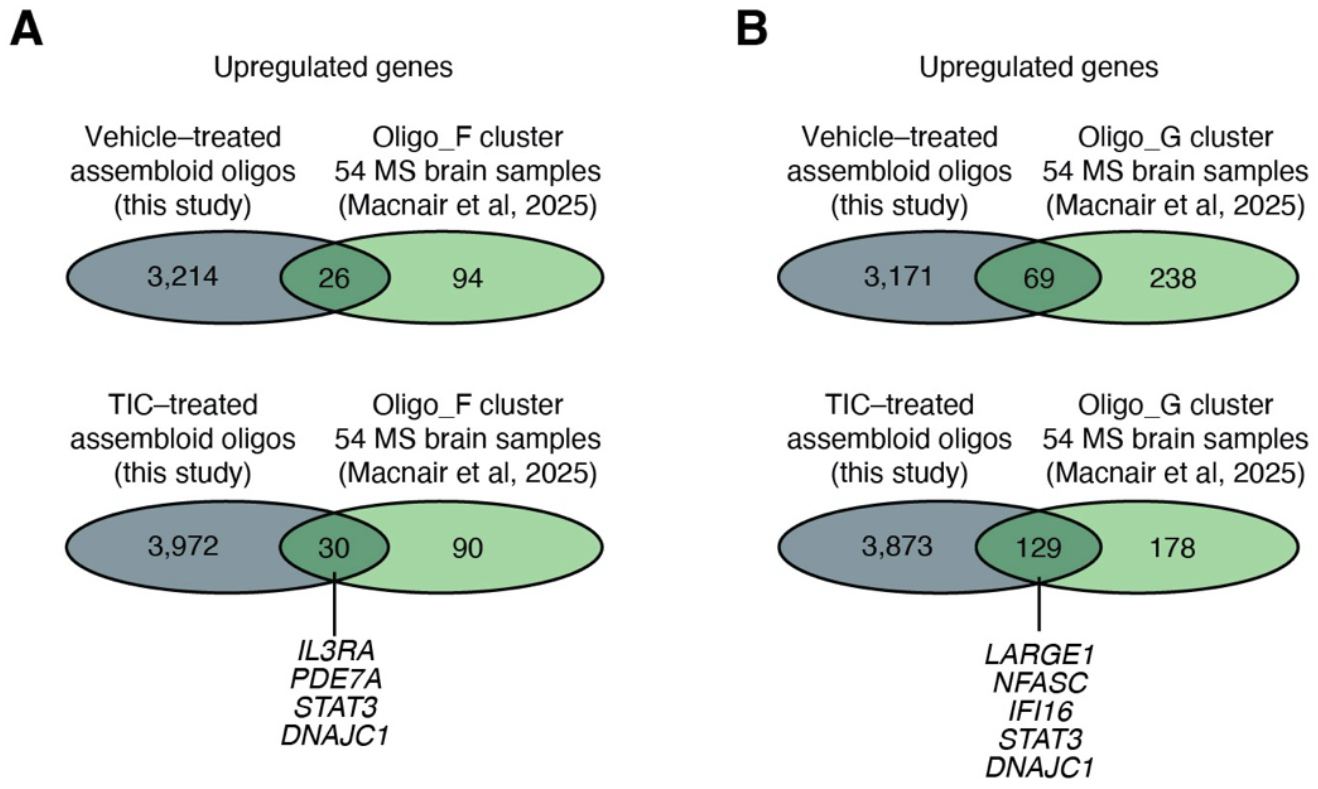
Overlap between cytokine-induced oligodendrocyte transcriptional changes and MS patient-derived disease signatures. **(A)** Venn diagrams representing comparison of differentially expressed genes in oligodendrocyte lineage cells from assembloids (left) treated with vehicle (top venn diagram) or TIC (bottom venn diagram) with significant markers of disease-associated MS oligodendrocyte cluster Oligo_F (right) from Macnair et al, 2025. Center number in the overlap represents number of shared genes, with arrow showing example genes in common with TIC-treated oligodendrocytes. All genes have adjusted p-value or false discovery rate (FDR) less than 0.05 and greater than 0.25 log_2_ fold change over their respective control. **(B)** Venn diagrams representing comparison of differentially expressed genes in oligodendrocyte lineage cells from assembloids (left) treated with vehicle (top venn diagram) or TIC (bottom venn diagram) with significant markers of disease-associated MS oligodendrocyte cluster Oligo_G (right) from Macnair et al, 2025. Center number in the overlap represents number of shared genes, with arrow showing example genes in common with TIC-treated oligodendrocytes. All genes have adjusted p-value or false discovery rate (FDR) less than 0.05 and greater than 0.25 log_2_ fold change over their respective control. See **Figure 4D** for pie chart comparison with TIC-treated oligodendrocytes from assembloids.

**Figure S6.**
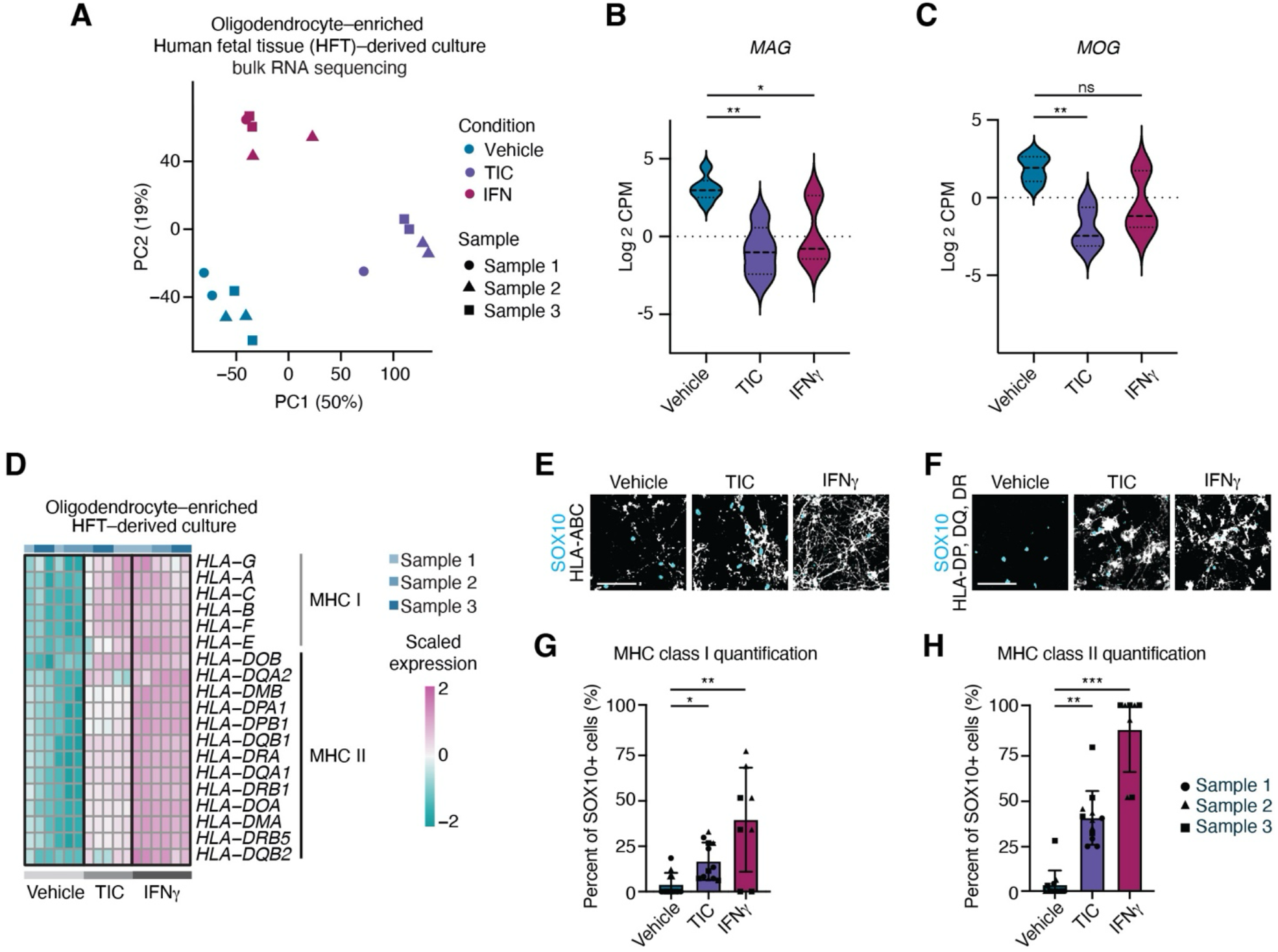
Validation of cytokine-induced changes in primary tissue. **(A)** Principal Component Analysis (PCA) plot showing oligodendrocyte-enriched cultures isolated from human primary fetal tissue showing PC1 and PC2 and colored by treatment: vehicle, TIC, or IFN-γ (n = 3 samples GW18–GW20, with 1–2 replicates per sample per condition). See **Figure S4A** for immunopanning scheme, and **Figure S4B, C** for enrichment validation. **(B)** Violin plot showing Log_2_CPM gene expression of oligodendrocyte marker *MAG* in vehicle, TIC, or IFN-γ treated oligodendrocyte-enriched cultures derived from human primary fetal tissue. Lines in violins represent median and first and third quartile. Mean ± SEM: Vehicle = 3.07± 0.33, TIC = -0.95 ± 0.72, IFN-γ = 0.13 ± 0.82; n = 3 samples GW18–GW20, with 1–2 replicates per sample per condition. Differential expression analysis was performed using limma-voom with empirical Bayes moderation on Log_2_CPM– transformed data; p-values were adjusted using the Benjamini–Hochberg method first, then doubled as a further Bonferroni adjustment for multiple comparison. *MAG*: TIC vs Vehicle p = 0.002, IFN-γ vs Vehicle p = 0.01. Statistical significance is denoted as p < 0.05 (*) and p < 0.01 (**). **(C)** Violin plot showing Log_2_CPM gene expression of oligodendrocyte marker *MOG* in vehicle, TIC, or IFN-γ treated oligodendrocyte-enriched cultures derived from human primary fetal tissue. Lines in violins represent median and first and third quartile. Mean ± SEM: Vehicle = 1.85± 0.35, TIC = -1.97 ± 0.59, IFN-γ = -0.47 ± 0.78; n = 3 samples GW18–GW20, with 1–2 replicates per sample per condition. Differential expression analysis was performed using limma-voom with empirical Bayes moderation on Log_2_CPM– transformed data; p-values were adjusted using the Benjamini–Hochberg method first, then doubled as a further Bonferroni adjustment for multiple comparison. *MOG*: TIC vs Vehicle p = 0.004, IFN-γ vs Vehicle p = 0.05. Statistical significance is denoted as p < 0.01 (**). **(D)** Heatmap showing MHC Class I and II expression from bulk RNAseq data in vehicle, TIC, or IFN-γ treated oligodendrocyte-enriched fetal culture (n = 3 samples, with 1–2 replicates per sample per condition). Z-scores were computed within each gene across conditions. **(E)** Representative images of each condition with immunohistochemistry of SOX10 and MHC Class I (HLA-ABC). **(F)** Representative images of each condition with immunohistochemistry of SOX10 and MHC Class II (HLA-DP, DQ, DR). **(G)** Quantification showing the percent of SOX10+ cells expressing MHC Class I in vehicle, TIC, or IFN-γ treated oligodendrocyte-enriched cultures. Vehicle = 3.8% ± 1.8%, TIC = 16.3% ± 2.9%, IFN-γ = 38.5% ± 9.8%; n = 2-3 samples GW18–GW20, 1-2 wells per sample, 2 images per well; Kruskal-Wallis test with Dunn’s multiple comparisons: vehicle vs TIC p = 0.034, vehicle vs IFN-γ p = 0.002). Statistical significance is denoted as p < 0.05 (*) and p < 0.01 (**). **(H)** Quantification of percent of SOX10+ cells expressing MHC Class II with vehicle, TIC, or IFN-γ treatment. Vehicle = 3.3% ± 2.2%, TIC = 39.3% ± 4.1%, IFN-γ = 86.5% ± 8.0%; n = 2-3 samples GW18– GW20, 1-2 wells per sample, 2 images per well; Kruskal-Wallis test with Dunn’s multiple comparisons: vehicle vs TIC p = 0.003, vehicle vs IFN-γ p < 0.0001). Statistical significance is denoted as p < 0.01 (**) and p < 0.001 (***). Scale bar: 100 μm (**E**), 100 μm (**F**).

**<u>Table S1.</u> Cell line information, reagents, and differential gene expression datasets.**

This multi-sheet Excel file includes detailed lists of cell lines used in each experiment, reagents, and differential gene expression lists. The first sheet is a table of contents with a description for each sheet of the file (**Table S1A–M**).

